# Lenacapavir-induced Lattice Hyperstabilization is Central to HIV-1 Capsid Failure at the Nuclear Pore Complex and in the Cytoplasm

**DOI:** 10.1101/2025.04.18.649524

**Authors:** Arpa Hudait, Ryan C. Burdick, Ellie K. Bare, Vinay K. Pathak, Gregory A. Voth

## Abstract

Lenacapavir (LEN) is the first HIV-1 capsid inhibitor approved for clinical use. It inhibits multiple steps of the viral life cycle; however, the molecular details of the effect of LEN on capsid structure and the mechanistic steps of the inhibition are not understood. Recent studies show that intact cone-shaped capsids and capsids with LEN-induced breaks can dock at nuclear pore complexes (NPC), but only intact capsids enter the nucleus. In this work, we combined large-scale coarse-grained molecular dynamics simulations and live-cell imaging to investigate the stepwise mechanism of docking of LEN-treated capsids into the NPC. Capsids bound to substoichiometric concentrations of LEN can reach the NPC central channel. As the capsid advances to the nuclear end, lattice defects are formed at the pentamer-hexamer interface – primarily at the narrower end – leading to pentamer dissociation. Dissociation of pentamers is detrimental to capsid integrity, leading to both rupture of the narrow end and destabilization of the hexamer-hexamer interface. Structural analysis of LEN-capsid complexes in our simulations demonstrates heterogeneous hyperstabilization and loss of the essential pliability of the capsid protein lattice. Live-cell imaging of HIV-1 cores labeled with two different fluorescent markers showed that LEN-treated ruptured capsids were docked at the NPC but were not imported into the nucleus. We conclude that LEN contributes to the loss of capsid elasticity and integrity, inhibiting HIV-1 nuclear entry and replication. Our findings demonstrate that altering viral material properties can be an effective strategy for designing antiviral drugs that disrupt viral core nuclear entry.

## Introduction

After human immunodeficiency virus type 1 (HIV-1) enters the cell, several distinct steps precede the integration of the viral genome into the host genome (1, 2). HIV-1 capsid consists of ∼250 capsid (CA) protein hexamers and 12 pentamers to form a cone-shaped structure (3-6). This capsid structure is commonly referred to as an HIV-1 “core”. The interior of the capsid contains the viral genome and essential replicative enzymes and is the site of reverse transcription. Nuclear pore complexes (NPC) are multiprotein assemblies embedded in the nuclear envelope that regulate nucleocytoplasmic transport of host proteins, as well as the passage of HIV-1 capsid into the nucleus (7). Early mechanistic models had proposed that viral cores uncoat in the cytoplasm or at the NPC central channel since the size of the capsid either exceeds or is comparable to the diameter of the NPC central channel (8-11). In contrast, recent studies indicate that viral cores remain intact during nuclear entry and terminally disassemble near the integration sites <1.5 hours before integration (12, 13). Consistent with these results, cryo-electron tomography (cryo-ET) images have revealed that the NPC central channel dimension is amenable for docking and import of intact cone-shaped capsids into the nucleus (14, 15). Our recent computational modeling demonstrated that the entry of intact capsids to the NPC central channel is regulated by the shape and orientation of the approach (16). Specifically, cone-shaped capsids, when approaching from the narrow end, progressively dilate the central channel, which provides an energetically favorable pathway of import. The sequential steps that regulate the nuclear entry of the capsid are potential targets for CA inhibitors (17, 18).

Lenacapavir (LEN) is the first HIV-1 capsid inhibitor approved for clinical use in People with HIV (PWH) at both the early and late stages of HIV-1 infection (19). The cellular proteins nucleoporin (NUP) 153 and cleavage and polyadenylation specificity factor 6 (CPSF6) facilitate the transport of the capsid to the nuclear basket and then to the nuclear interior (20-23). These cellular proteins are rich in phenylalanine-glycine (FG) motifs, which bind to a CA hydrophobic FG-binding pocket (24, 25), to which LEN also binds (17). At the binding pocket, LEN makes extensive contacts with adjacent CA monomer residues mediated through electrostatic and Van der Waals interactions (17). Importantly, LEN is a more potent capsid inhibitor than PF3450074 (PF74), another capsid binding inhibitor, and at much lower concentrations (26). Recent studies have demonstrated that LEN treatment can lead to loss of capsid integrity(26, 27). Furthermore, in the presence of LEN, there is an increase in the number of viral cores in the cytoplasm (17). This raises the question of whether LEN treatment can modulate the integrity of the capsid during nuclear entry.

The human NPC is a ∼120 MDa macromolecular assembly consisting of 30 distinct NUPs (7). These NUPs form multiple heterooligomeric complexes, which assemble into a stacked outer cytoplasmic ring (CR), nuclear ring (NR), and inner ring (IR). The CR and NR consist of eight heterooligomeric Y-complex dimers arranged in a head-to-tail arrangement. The IR consists of eight spokes that are connected through linkers that provide conformational flexibility (28). The disordered NUPs, rich in capsid-binding FG dipeptide motifs, are also bound to the IR spokes, forming a cohesive network at the central channel (29). FG-NUPs are key to the initial docking and translocation of the viral capsid (30). The mechanism of capsid translocation driven by interaction with FG-NUPs is comparable to the transport of large cargoes mediated by transporters in the karyopherin family (31). All-atom molecular dynamics (AA MD) simulations of membrane-embedded NPC to investigate the dynamics of capsid docking would require over a billion atoms and are computationally infeasible due to the system size and time scales of the processes of interest. Coarse-grained (CG) MD simulations, however, performed with “bottom-up” CG models can afford important mechanistic insight into the dynamics of these cellular processes. “Bottom-up” CG models are systematically derived from the underlying atomistic-level interactions to reproduce the molecular behavior as projected to a coarser representation (32).

In contrast, “top-down” CG models have also been used for modeling nuclear entry (33) and capsid elasticity (34). While the CG simulations of the latter are argued by those authors to connect with atomic force microscopy (AFM) experiments, they do not incorporate solvent effects or the ribonucleoprotein (RNP) and they do not include LEN nor the NPC. Capsid strain and deformations have also been investigated previously using AAMD (35) with explicit IP6 and RNP. To efficiently simulate the nuclear entry of HIV-1 capsid, we developed a “bottom-up” CG model (16) of the human NPC and HIV-1 capsid from high-resolution cryo-ET structures (7). Our CG MD simulations of capsid docking at the NPC central channel demonstrated that lattice elasticity and the flexibility of the pore are key for the passage of the intact capsid through the channel. Importantly, the computational predictions of capsid docking to the NPC central channel have been recently validated in a HIV-1 core import at the NPC using cryo-ET (36), demonstrating how systematically derived “bottom-up” CG models can accurately predict molecular details of complex biomolecular processes. To our knowledge, a comprehensive study demonstrating the stepwise details of the effects of LEN on a biologically realistic capsid (that harbors a model for the RNP) during nuclear docking (i.e. traversal from the cytoplasmic side to the central channel) has been lacking.

In this work, we investigate the docking of LEN-bound cone-shaped capsids into the NPC central channel using a combined computational and experimental approach. Our goal is to shed light on the interplay between LEN, capsid, and FG nucleoporins at the NPC that modulates capsid structural integrity. In our CG MD simulations, capsids bound to substoichiometric concentrations of LEN successfully dock into the NPC central channel. However, as the capsid approaches the nuclear end of the channel when translocating through the FG repeat permeability barrier, defects are formed at the hexamer-pentamer interface, predominantly at the narrow end. Computational structural analysis of LEN-bound capsid cones reveals hyperstabilization and stiffening of the hexameric lattice since LEN can only bind to the CA hexamers. The formation of the defects and lattice hyperstabilization leads to the processive loss of pentamers, followed by destabilization of the hexamer-hexamer interface, resulting in capsid rupture. We also observe a similar mechanism of stepwise rupture of the freely diffusing LEN-capsid complex, which emulates the capsid in cytoplasmic environment. Consistent with our CG MD simulations, live-cell imaging of HIV-1 cores labeled with two different fluorescent markers in cells reveals that LEN treatment caused HIV-1 cores docked at the NPC to rupture and that the broken capsids remained bound to the NPC, but the capsid contents are lost. Our results demonstrate that stress from NPC can accelerate the rupture of LEN-treated hyperstabilized capsids relative freely diffusing capsids not docked at the NPC.

## Results

### Coarse-grained modeling and simulation

We recently developed a “bottom-up” solvent-free membrane-embedded CG model of the human NPC that consists of 8 copies of the dimerized Y-complex and IR spokes (16). The membrane-embedded composite NPC model is shown in **Fig. 1A**. The FG-rich NUP62 is modeled as heterotrimeric subcomplex along with NUP54 and NUP58. The disordered NUP98 containing capsid-binding FG-motifs are tethered to the NPC central and create a hydrogel-like environment through multivalent interactions (29, 37, 38). We used inter-NUP98 associative interactions that result in extended conformations of the NUP98 chains, forming a mesh-like environment at the channel and closely mimicking the NPC environment *in vivo* (37). Importantly, we previously demonstrated that capsid docking is promoted by the extended conformations of NUP98 chains (16). As the capsid translocates to the nuclear side, the conformation state of NUP98 chains in the mesh remains mostly unaltered (i.e., the extended state is preserved, **Fig. S1**). **Movie S1** depicts the capsid cone (approaching from the narrow end) docking at the NPC central channel.

**Figure 1.**
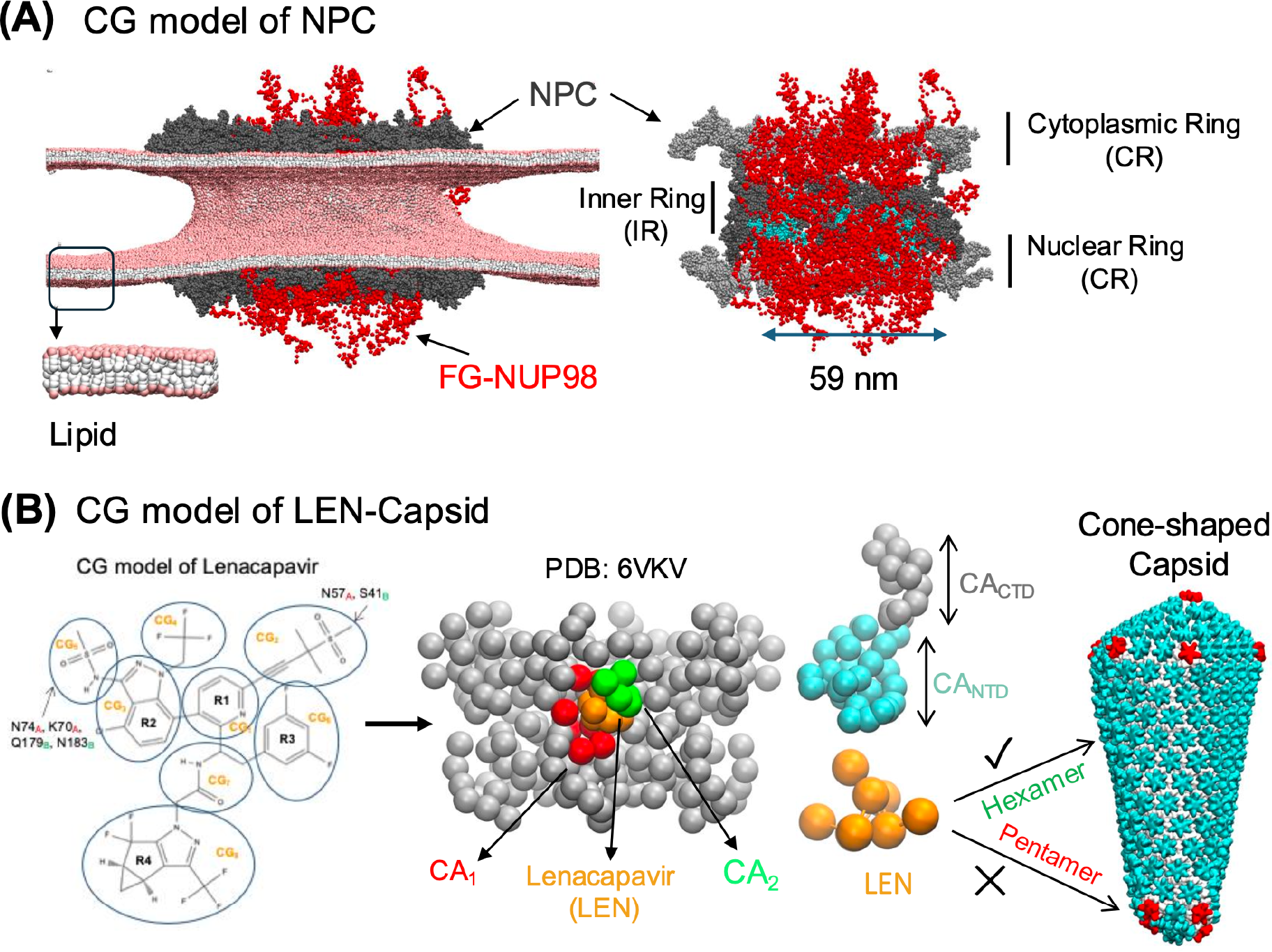
Overview of the CG molecular model of the NPC, HIV-1 capsid, and LEN. **(A)** The left panel shows the composite membrane-embedded CG model of human NPC. The NPC is shown in gray spheres. The disordered NUP98 chains are shown in red chains. The nuclear membrane is modeled with a 4-site CG lipid bilayer model. The CG bead representing the lipid headgroups are shown in pink spheres, interfacial and tail beads are shown in white spheres. In the right panel, the cross-section of the NPC is shown. The CR and NR are shown in silver to distinguish from the IR (shown in gray spheres). The FG-NUP62 is modeled as a heterotrimeric subcomplex along with NUP54 and NUP58 is shown in cyan spheres (occluded by the FG-NUP98 mesh). The diameter of the central channel is 59 nm. **(B)** The left panel shows the CG representation of LEN with CG sites 1-8 labeled on the LEN chemical representation. The center panel shows the CG-mapped representation of LEN bound to a CA hexamer. The CG-mapped structure was generated from the X-ray crystal structure PDB: 6VKV. A CG LEN molecule is shown in orange beads. The CG sites of adjoining CA monomers CA_1_ and CA_2_ that are in contact with LEN are shown in red and green, respectively. In the right panel, the CG representation of the CA monomer is shown. The CA_NTD_ and CA_CTD_ domains are shown in cyan and silver spheres, respectively. In the snapshot of the cone-shaped capsid, the CA_NTD_ of the hexamer is shown in cyan, and the pentamer is shown in red. The CA_CTD_ domain of both hexamer and pentamer is shown in silver.

It should also be noted that in these and in our past CG MD simulations (16) of capsid docking into the NPC, it is infrequently observed that the NPC ring can “break” and the capsid docks in a somewhat sideways orientation (Fig. S2). This behavior has been recently suggested experimentally as well (33), though we considered it to be a minor component of the ensemble of capsid-NPC docking events. We also note that in our CG MD model simulations, we do not need to apply a force to observe the capsid docking behavior (it occurs as an electrostatic “ratchet”), which may be contrasted with a different CG simulation approach to this docking behavior based on the top-down Martini CG model (33).

Lenacapavir (**Fig. 1B**) is modeled with a 8-site CG model directly from the high-resolution X-ray crystal structure of LEN complexed to CA hexamer (PDB: 6VKV) (17). The CA-LEN non-bonded associative interactions (electrostatic and Van der Waals) are modeled with attractive interactions of distinct energy scales. In our CG model, stronger attractive interactions are used to model the CA-LEN hydrogen bonding interactions relative to the van der Waals interactions. The magnitude of the attractive interactions was adjusted to capture the substoichiometric binding of LEN to CA hexamers (26). We model LEN and CA interactions such that LEN molecules can only bind to CA hexamers, and all interactions to CA pentamers are turned off, as in experiments, CA selectively associates with hexamers (25, 39). We simulated LEN binding to the capsid cone (in the absence of NPC), which resulted in a substoichiometric binding (∼1.5 LEN per CA hexamer), consistent with experimental data (40). Using the aforementioned CG models, we performed CG MD simulations of LEN-capsid docking at the NPC central channel. The LEN-capsid complex (bound stoichiometry of ∼1.5 LEN per CA hexamer) was placed at the cytoplasmic side of the NPC, with the narrow end pointing to the central channel, and touching the cytoplasmic end of the central channel. Complete details of the simulation setup are provided in the *SI Appendix Methods*.

### Competition between FG-NUP98 and LEN binding regulates the early stages of the capsid docking into the NPC central channel

To simulate capsid docking into the NPC central channel, we placed LEN-treated cone-shaped capsids at the cytoplasmic end of the NPC (coplanar to the cytoplasmic Y-complex). To model the ribonucleoprotein (RNP) complex, we added two polymeric chains in the capsid interior. The RNP complex minimally emulates the effect of nucleocapsid (NC) and RNA chains (the model details in *SI Appendix Methods*). We performed two independent replicate simulations. The simulations were performed for 200 × 10^6^ CG MD timesteps (τ_*CG*_ = 50 fs). From these simulations, to characterize the capsid docking dynamics, LEN, and NUP98 binding to the capsid, we calculated the time series statistics of three following metrics: 1. distance (*D*_*Cap-NPC*_) between the geometric center of the capsid and equatorial midplane of the NPC inner ring along the channel axis; 2. the number of FG sites of NUP98 (*f*_*FG-NUP98*_) directly in contact with the CA monomers; and 3. the number of LEN molecules bound to the capsid (*N*_*LEN*_). The number of FG sites of NUP98 in contact with CA monomers are calculated as normalized by the number of CA monomers of all the hexamers. We plot the time series in Fig. 2 of the LEN-capsid complex until the narrow end of the capsid reaches the nuclear end of the NPC, prior to the rupture events of the capsid.

**Figure 2.**
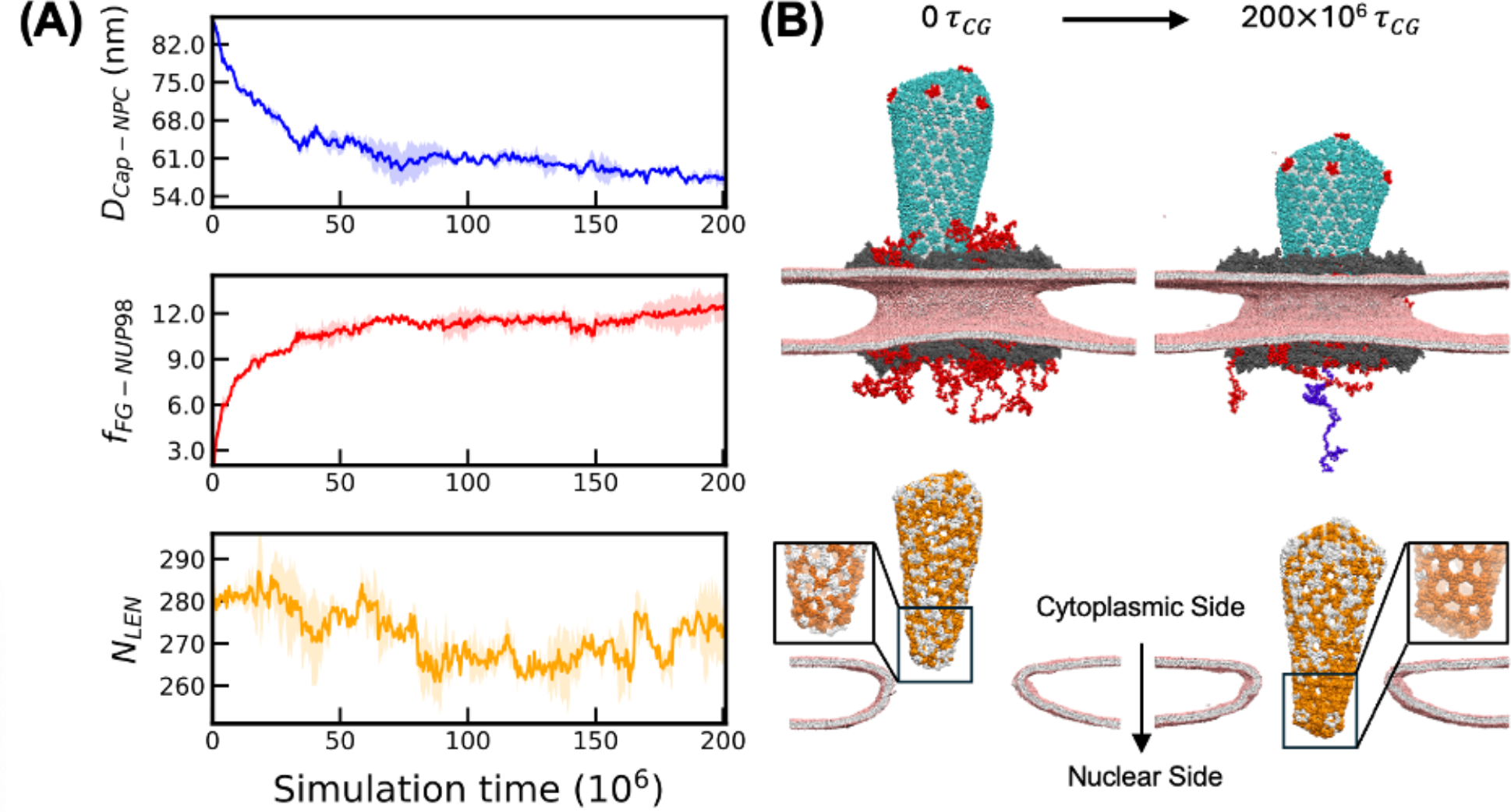
Competition between NUP98 and LEN during docking of LEN-treated capsid. **(A)** The time series of the distance (*D*_*Cap-NPC*_) between the geometric center of the LEN-bound capsid and equatorial midplane of the NPC inner ring along the channel axis (upper panel), the number of FG sites of NUP98 (*f*_*FG-NUP98*_) directly in contact with the CA monomers (middle panel), and the number of LEN molecules bound to the capsid (*N*_*LEN*_) (lower panel) are plotted. Here, *f*_*FG-NUP98*_ denotes the total number of FG sites that are in the vicinity of the CA hydrophobic FG-binding pocket (within 4 nm radius). **(B)** The upper panel shows the initial configuration and the final configuration of the capsid at the end of 200 × 10^6^ τ_*CG*_. In the snapshot at 200 × 10^6^ τ_*CG*_, one RNP chain (blue spheres) partly extrudes out concomitant to the first appearance of defects at the pentamer-hexamer interface. The color scheme is the same as in Fig. 1. In the lower panel, only the CTD domain is shown for each CA monomer. The CA monomers to which at least one LEN molecule is bound are shown in white spheres. The CA monomers to which no LEN molecule is bound are shown in orange spheres. Note the high density of orange spheres located at the narrow end of the capsid in the right panel (docked capsid). This indicates that some LEN molecules are displaced by the NUP98 mesh to allow capsid docking. A cutaway view of the nuclear membrane is also shown to represent the degree of capsid docking from the cytoplasmic side toward the nuclear side. The NPC, LEN, and NUP98 are not shown for figure clarity. For all the plots, the solid lines are the mean values calculated from the time series of two independent replicas, and the shaded region is the standard deviation at each timestep.

Note that in this work, we use the term “capsid docking” instead of “capsid translocation”. The process of capsid docking denotes the association of the capsid to the NPC central channel. In contrast, capsid translocation denotes the entry of the capsid into the nuclear basket, followed by to the nuclear interior. The simulation of the capsid translocation to the nuclear end requires a structural model of the nuclear basket. However, when we performed this study, a structural model of the human nuclear basket was not available.

The time series profiles of docking of the LEN-capsid complex from the cytoplasmic to the nuclear side of the NPC reveal key insight into the dynamics of capsid docking (**Fig. 2A**). Initially, the LEN-capsid complex undergoes rapid passage as the number of FG sites in contact with CA steeply increases. The kinetics of passage gradually decrease as the central channel encounters the wide regions of the capsid, and there is an increasing degree of steric interaction between the capsid and the non-FG-NUPs. At this stage, the capsid is bound to the NPC mediated via the mesh-like network created by the NUP98 chains. The continuing movement of the capsid towards the nuclear end at this stage will require disruption of the FG-mesh, dissociation, and formation of FG-CA contacts. We then compared the time series of the number of LEN molecules bound to capsid and FG-CA contacts. At the late stages of LEN-capsid docking, we observe competition between LEN binding and FG-CA interactions (**Fig. 2B**). A fraction of LEN molecules bound at the narrow end dissociate to allow NUP98 binding to the capsid (**Fig. 2B** and **Fig. S3**). In our simulations, LEN binding to the capsid is substoichiometric. It is possible that at significantly higher concentrations of LEN, NUP98 binding to the capsid will be adversely impacted. Therefore, our modeling demonstrates that at significantly high concentration LEN can inhibit the efficient binding of the viral cores to the NPC, resulting in an increased number of cores in the cytoplasm.

### Distinct lattice defects at the pentamer-hexamer interface at the narrow end culminate in rupture

Visualization of the NPC docking trajectories of LEN-capsid complexes revealed several key steps that lead to the loss of core integrity (**Fig. 3A, SI Movie 2 and 3**). In our simulations, pentamer-hexamer contacts are transiently disrupted, leading to dissociation of the CA pentamer rings both at the narrow and wide ends. CA pentamers at the narrow end are likely to dissociate faster than at the wide end, likely a result of weakened contacts between hexamers and pentamers due to higher curvature. Molecular fluctuations result in the loss of pentamer-hexamer contacts and lead to irreversible dissociation of the pentamers from both the narrow and the wide end (**Fig. 3B**). Concurrent with the appearance of defects and dissociation of pentamers, we also observe RNP complex extruding through the lattice defects. Dissociation of multiple pentamers from the narrow end is detrimental to capsid integrity, resulting in lattice rupture. After rupture of the lattice, we observe the formation of defects at the hexamer-hexamer interface away from the narrow end, demonstrating further weakening of the lattice integrity. Although high computational cost precluded us from continuing these CG MD simulations, we expect these defects at the hexamer-hexamer interface to propagate from the high curvature to low curvature end of the capsid.

**Figure 3.**
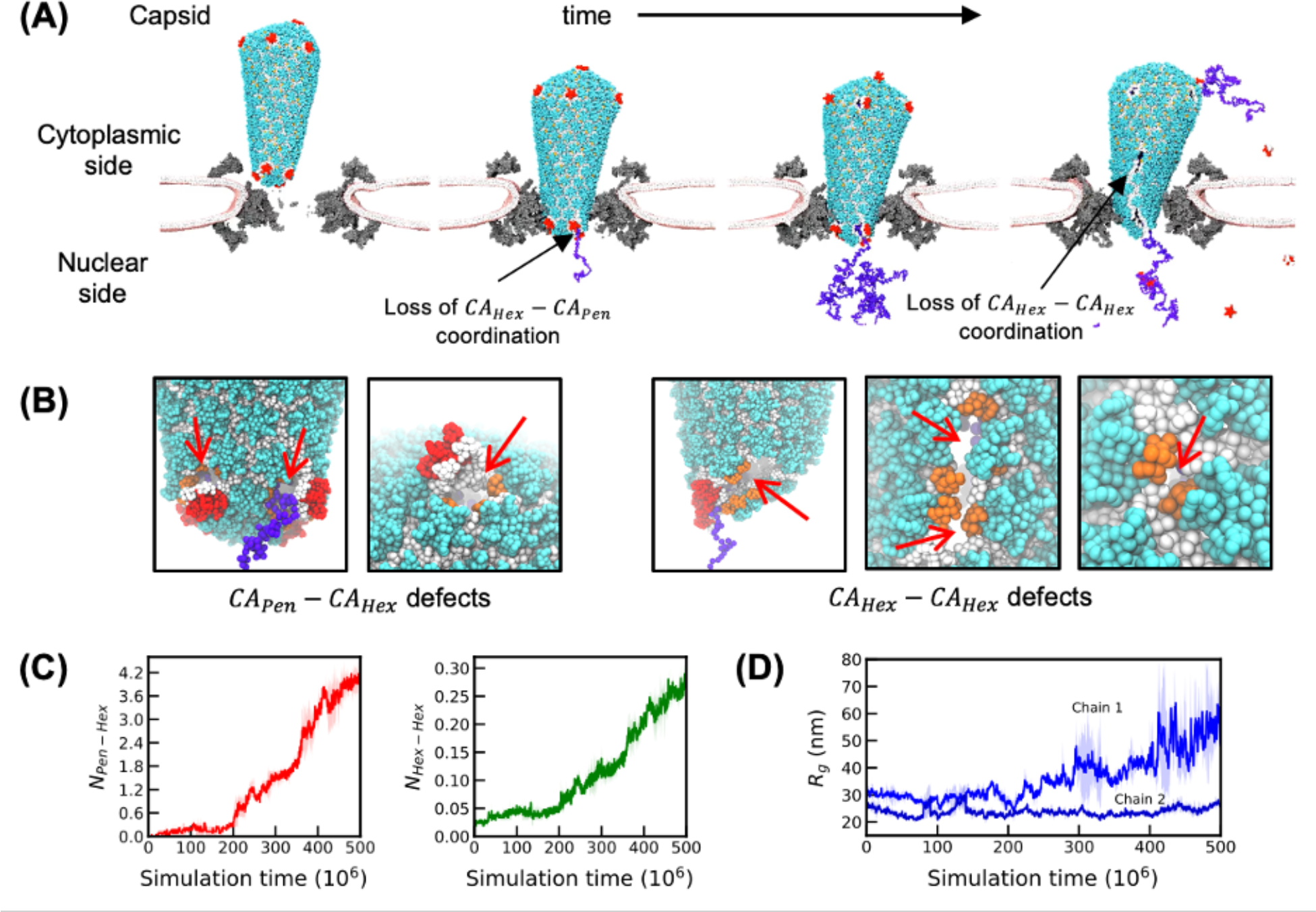
Stepwise rupture of capsid treated with LEN during NPC docking. **(A)** The NTD domain of CA hexamer and pentamer are shown in cyan and red spheres, respectively. The CTD domain of all CA monomers is shown in white spheres. The RNP chains are shown in blue. The NPC (cutaway view) is shown in gray, and the nuclear membrane is shown in pink (head group) and white (other groups). The initial events of CA-CA contact disruption at the hexamer-pentamer interface (*CA*_*Hex*_ − *CA*_*Pen*_) at the narrow end are labeled with arrow. In the leftmost snapshot, the RNA extrudes out of the defect as a result of partial dissociation of the pentamer. In the final stage, nucleation of cracks occurs at the hexamer-hexamer interface (*CA*_*Hex*_ − *CA*_*Hex*_) and extends from the narrow to the wide tip. **(B)** Snapshots of the defects at the hexamer-pentamer interface and hexamer-hexamer interface. CA_CTD_ domains of the hexamers that constitute the defect site are highlighted in dark orange spheres. Note that the CG beads constituting the CA_CTD_ domains at the defect site are disconnected from at least one nearest neighbor, leading to the locally cracked lattice (marked with red arrows). **(C)** Time series of the degree of defects at the pentamer-hexamer (*N*_*Pen*-*Hex*_) and hexamer-hexamer (*N*_*Hex*-*Hex*_). Note that in *N*_*Pen*-*Hex*_ and *N*_*Hex*-*Hex*_ are calculated by normalizing by total number of CA pentamer (12) and hexamer rings (209) respectively. The left panel shows the time series of the undercoordinated CA monomers at the pentamer-hexamer interface. The right panel shows the time series of the undercoordinated CA monomers at the hexamer-hexamer interface. **(D)** Time series of the radius of gyration (*R*_*g*_) of the two RNP chains. The solid lines are the mean values calculated from the time series of two independent replicas, and the shaded region is the standard deviation at each timestep.

To characterize the molecular details of lattice rupture, we estimated the number of CA monomers that are under-coordinated (defined as the monomers with less than three neighbors at the trimeric CTD-CTD interface). Furthermore, we denoted defects as under-coordinated CA monomers of the hexamers at the pentamer-hexamer and hexamer-hexamer interface as *CA*_*Pen*_ − *CA*_*Hex*_ and *CA*_*Hex*_ − *CA*_*Hex*_, respectively (**Fig. 3B**). In addition, we represented the defects as the number of under-coordinated CA monomers of the hexamers at the pentamer-hexamer-pentamer and hexamer-hexamer interface as *N*_*Pen*-*Hex*_ and *N*_*Hex*-*Hex*_ (**Fig. 3C**). The time series profile of *N*_*Pen*-*Hex*_ and *N*_*Hex*-*Hex*_ indicates that these defects appear concurrently (∼200 × 10^6^ τ_*CG*_). To characterize the correlation between the formation of lattice defects and RNP conformation, we calculate the radius of gyration (*R*_*g*_). The time series of *R*_*g*_ (**Fig. 3D**) demonstrates that one RNP chain begins to extrude from the lattice defects at the narrow end coincident with the first appearance of a lattice defect (∼200 × 10^6^ τ_*CG*_). The second RNP chain remained associated to the CA_CTD_ during our simulations.

To summarize, our CG MD simulations demonstrate that LEN-treated cores can efficiently dock at the NPC central channel, which is followed by the appearance of defects at the narrow and wide ends. These defects lead to dissociation of CA pentamers, crack formation at the hexamer lattice, and partial release of the RNP complex. Our results thus suggest that cores bound to substoichiometic concentrations of LEN efficiently dock at the NPC and then first rupture at the narrow end when bound to the NPC central channel.

### Ruptured LEN-viral complexes remain bound to the NPC

We developed an experimental live-cell imaging assay (details in *SI Appendix Methods*) to assess the impact of LEN on viral cores that are stably docked at NPCs. To visualize viral cores, we labeled the capsid lattice with GFP-CA as previously described (12) and a capsid fluid phase content with cmHALO bound to JF646 fluorescent dye, which was similar to previously described GFP content marker labeling (13) **(Fig. 4A; Fig. S4A-E)**. This dual-labeling strategy allowed us to distinguish intact GFP-CA-labeled viral cores, which retain cmHALO, from broken ones, which have lost cmHALO. To visualize NPCs, we generated a HeLa cell line stably expressing the nuclear pore protein POM121 fused to HALO, labeled with JF549 dye. Time-lapse imaging began 2 hours post-infection, with either DMSO (control) or LEN added after the first frame **(Fig. S4F)**. We identified GFP-CA-labeled viral cores that remained stably docked at the nuclear envelope throughout the 15-minute observation period (**Fig. 4B**). Few viral cores remained at the nuclear envelope in either treatment condition (**Fig. 4C**), aligning with our previous finding that most cytoplasmic viral cores fail to stably associate with the nuclear envelope (10). At the start of imaging, most of the stably docked viral cores retained cmHALO (83%-89%; **Fig. 4D**), indicating they were intact. However, treatment with 100 nM LEN led to a rapid loss of cmHALO in nearly all viral cores compared to DMSO-treatment (97% vs. 8%, respectively; **Fig. 4E**), occurring approximately 4 minutes after LEN addition (**Fig. 4F**). In agreement with the CG MD simulations, these findings suggest that LEN induces rupture of NPC-docked viral cores and that the broken capsid lattice remains associated with the NPC.

**Figure 4.**
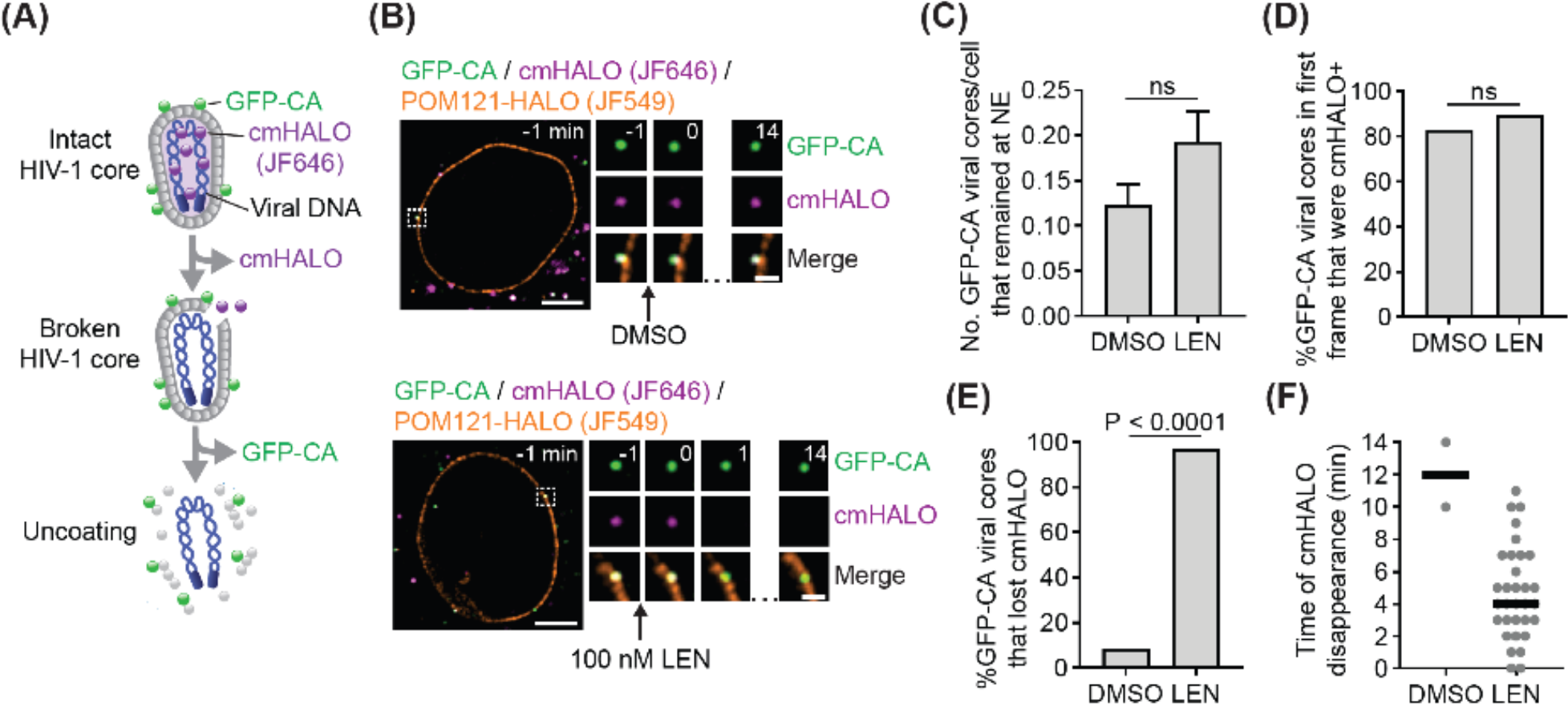
Live-cell imaging of LEN-induced rupture of HIV-1 cores docked at nuclear envelope. **(A)** Dual-labeled viral cores: the capsid lattice is labeled with GFP-CA, and the capsid content is labeled with content marker HALO (cmHALO; JF646 dye). **(B)** Representative GFP-CA labeled viral core in a cell expressing POM121-HALO (JF549 dye) that was stably associated with the nuclear envelope, retaining cmHALO in a DMSO-treated control cell (top) or losing cmHALO within 1 min of 100 nM LEN addition (bottom). **(C)** Number of GFP-CA labeled viral cores per cell that remained at the nuclear envelope during the 15-minute observation period. A total of 237 DMSO-treated cells and 192 LEN-treated cells were analyzed. **(D)** Percentage of GFP-CA labeled viral cores that were cmHALO^+^ at the start of imaging. GFP-CA labeled viral cores were analyzed for DMSO-treated (29 total) and nm LEN-treated (37 total) cells. **(E)** Percentage of GFP-CA labeled viral cores that lost cmHALO during the 15-minute observation period. **(F)** Time (minutes) of cmHALO disappearance following DMSO or LEN addition. *P* values were calculated using Fisher’s exact test; ns, not significant (*P* > 0.05).

### Rupture of LEN-treated freely diffusing capsids exhibits slower kinetics relative to NPC-docked capsids in simulations

To investigate how LEN binding modulates capsid molecular features, we simulated with CG MD freely diffusing capsid cones (not bound to the NPC), with the initial unbound concentration of LEN amounting to a stoichiometry (LEN:CA) of ∼2.9:6. Here, the freely diffusing LEN-capsid complex minimally emulates the capsid at the cytoplasm. In the initial configuration, no LEN molecules were bound to the capsid. We started with a substoichiometric excess of unbound LEN molecules in the system to achieve stoichiometric saturation within the limited (relative to experiments) simulation timescales. We evolved the system for 500 × 10^6^ τ_*CG*_ and achieved a LEN-bound stoichiometry of ∼1.5:6. During these simulations, CA pentamer-hexamer contacts are only transiently broken. The number of defects progressively increases, albeit at a significantly slower rate compared to the NPC-docked capsids in the same CG simulation time (**Fig. S5**). In other words, fewer defects form in the free capsids compared to the NPC-docked capsids at similar time scales. Note that the number of LEN molecules bound to the capsid for the free capsid and NPC-docked capsids are nearly identical. Hence, the disparity in timescale of lattice rupture is not only because of the effect of LEN on capsid lattice properties. The disparity in the timescales of the appearance of lattice defects in NPC-bound and free capsid indicates that the confining stress and steric interactions at the central channel weaken the inter-CA ring interactions of the lattice. This can effectively decrease the barrier for dissociating the pentamer-hexamer contacts and facilitate lattice rupture. As a consequence, in the CG MD without sampling (see next paragraph) we observe lattice rupture in LEN-treated capsid cones that are docked at NPCs but not when they are free in CG MD simulations.

To observe rupturing of free capsids, we performed well-tempered metadynamics (WTMetaD) simulations (41). Briefly, WTMetaD is an accelerated sampling technique that can be used to explore high free energy barrier processes. In WTMetaD simulations, we used the mean coordination number (**Fig. S6**) between CA proteins in pentamers and in hexamers as the reaction coordinate (see *SI Appendix Methods*). For the WTMetaD simulations, we performed four replica simulations, each with a condensed and an uncondensed RNP complex. The two interacting polymeric chains are placed inside the capsid that emulates two 9000-nt RNP complexes, and modulating the interaction strength between the CG beads of the polymers controls the condensation state of the RNP (Details in *SI Appendix Methods*).

We previously demonstrated that the RNP complex inside the capsid contributes to internal mechanical strain on the lattice driven by CA_CTD_-RNP interactions and condensation state of RNP complex (16). The stepwise mechanism of lattice rupture for a free LEN-treated capsid observed in WTMetaD simulations is mostly similar to the NPC-bound capsid. First, pentamer-hexamer contacts are disrupted at the narrow end, followed by at the wide end. The progressive disruption of the pentamer-hexamer contacts is followed by cracks at the hexamer-hexamer interface, leading to rupture at the narrow end (**Fig. 5, SI Movies 4, 5**, and **Fig. S7**). The condensation state of the RNP provides significantly different molecular environments. The uncondensed RNP model occupies the entirety of the internal space of the capsid. In contrast, the condensed RNP remains localized at the wide end, stabilized by interactions with CA_CTD_.

**Figure 5.**
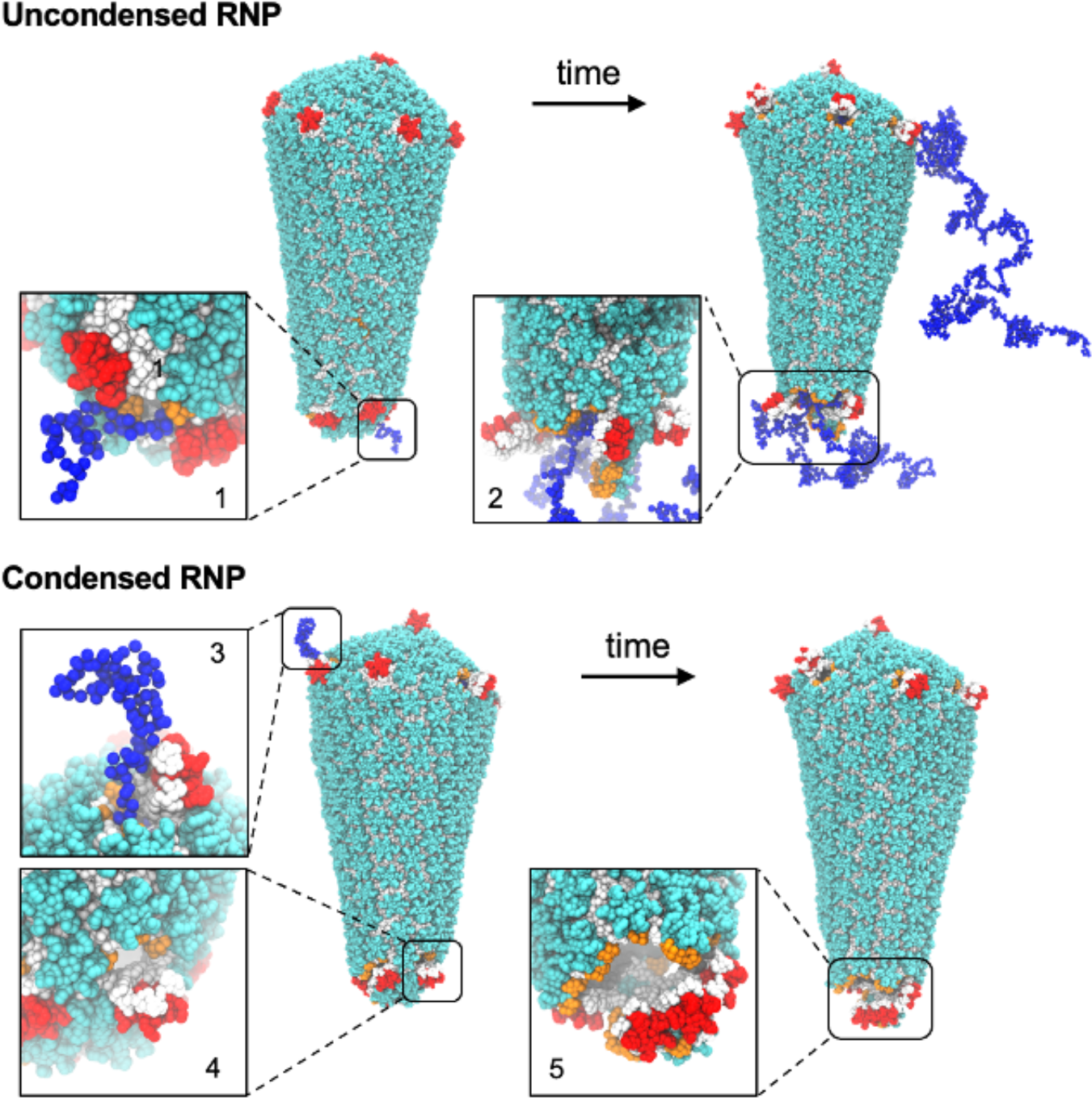
Molecular view of defects and LEN-induced rupture of free capsids. The color scheme of the capsid and RNP is the same as in Figure 3. CA_CTD_ domains of the hexamers that constitute the defect site are highlighted in orange spheres. The snapshots in the inset show zoomed-in view of representative defects. For the uncondensed RNP, snapshot 1 shows partial dissociation of pentamers at the narrow end, and RNP chains extrude out of the capsid interior. Snapshot 2 shows a ruptured narrow end. For the condensed RNP, snapshots 3 and 4 show defects arising from the partial dissociation of pentamers at the wide end and the narrow end. Condensed RNP localizes at the capsid wide end. In snapshot 3, the RNP extrudes out of the defect at the wide end. Finally, snapshot 5 shows rupture of the narrow end.

Based on the fluctuation of the reaction coordinate (mean coordination number between CA proteins in pentamers and in hexamers) in the WTMetaD simulations, we estimated the barrier of capsid disassembly to be ∼11 kcal/mol (**Fig. S8**). The free energy barrier for capsid dissociation for internal RNP in condensed and uncondensed state is nearly identical. Based on the free energy barrier of pentamer dissociation, we estimate the timescale of a single pentamer dissociation from the capsid to be ∼10 μs in CG simulation time. This can rationalize why we do not observe capsid rupture in unbiased simulations in the cytoplasm, and in contrast, capsid rupture is observed during NPC docking, where the rupture is facilitated by steric effects of the NPC central channel. In a recent experiment, the timescale of capsid rupture treated with 100 nM LEN is 4 min (42). A possible mechanistic pathway of capsid disassembly can be that multiple pentamers are dissociated from the capsid, and the remaining hexameric lattice remains stabilized by bound LEN molecules, before, the structural integrity of the remaining lattice is compromised.

Our results suggest that the condensation state of the RNP complex does not significantly impact the degree of lattice defects. This also raises the question as to whether the degree of reverse transcription in the capsid impacts the integrity of LEN-treated capsids. Both the NPC-docked and free capsid can have partially reverse transcribed DNA in them, which likely influences their condensation state. Future studies with detailed models of RNA-DNA conversion mimicking different stages of reverse transcription could be used to investigate how DNA compaction affects the degree of lattice defects.

### LEN binding to the capsid results in hyperstabilized lattice domains

To elucidate the molecular signatures regulating the correlation between structural heterogeneity of the capsid lattice and LEN binding, we characterized the distribution of LEN bound to the freely diffusing capsid from our simulations. We found that 1 or 2 LEN molecules are associated with CA hexamers at nearly equivalent probability (**Fig. S9**). We also observed infrequent association of 3 LEN molecules to the CA hexamer. This indicates that the probability of binding a third LEN molecule to a CA hexamer is impeded, likely due to steric effects that prevent the approach of an incoming molecule to a CA hexamer where 2 LEN molecules are already associated. Approximately 20% of CA hexamers remain unoccupied despite the availability of a large excess of unbound LEN molecules. This suggests a heterogeneity in the molecular environment of the capsid lattice for LEN binding.

The HIV-1 capsid undergoes volume fluctuations, leading to expansive and compressive strains that manifest into highly correlated striated patterns at the capsid lattice (16, 35). These striated patterns also demonstrate deviations from ideal lattice packing. We characterize the distortion of the capsid lattice using neighbor-averaged Steinhardt’s local bond order parameter (⟨*q*_6_⟩_*neigh*_) for each CA monomer (43, 44). Here, lower values of (⟨*q*_6_⟩_*neigh*_) are indicative of the distortion of the lattice contiguous to a CA monomer from ideal packing. We then identified in our simulation trajectories whether a LEN molecule binds to a CA monomer that is classified as part of an ordered (*LEN*_*o*_) or distorted (*LEN*_*d*_) lattice (**Fig. 6A**). We found that LEN molecules bind to the distorted CA lattice sites with higher propensity than the ordered lattice (approximately 2:1), demonstrating the heterogeneous nature of the CA-LEN interactions. To rationalize this disparity in binding propensity, we calculated the distribution of the potential energy of CA-LEN interactions in the LEN-capsid complex for the LEN molecules bound to the ordered or distorted lattice sites (**Fig. 6B**). The distribution demonstrates that the binding of LEN to the distorted lattice sites is energetically favorable. Since LEN localizes at the hydrophobic pocket between two adjoining CA monomers, it is sterically favorable to accommodate the incoming molecule at a distorted lattice site. This can likely be attributed to the higher available void volume at the distorted lattice relative to an ordered lattice, the latter being tightly packed (35). This also allows the drug molecule to avoid the multitude of unfavorable CA-LEN interactions and establish the energetically favorable interactions leading to a successful binding event. Here we denote unfavorable CA-LEN interactions as all interactions other than the electrostatic and van der Waal interactions that lead to CA-LEN binding (17). These results taken together point to the heterogeneity in the molecular environment of the capsid lattice for LEN binding.

**Fig. 6.**
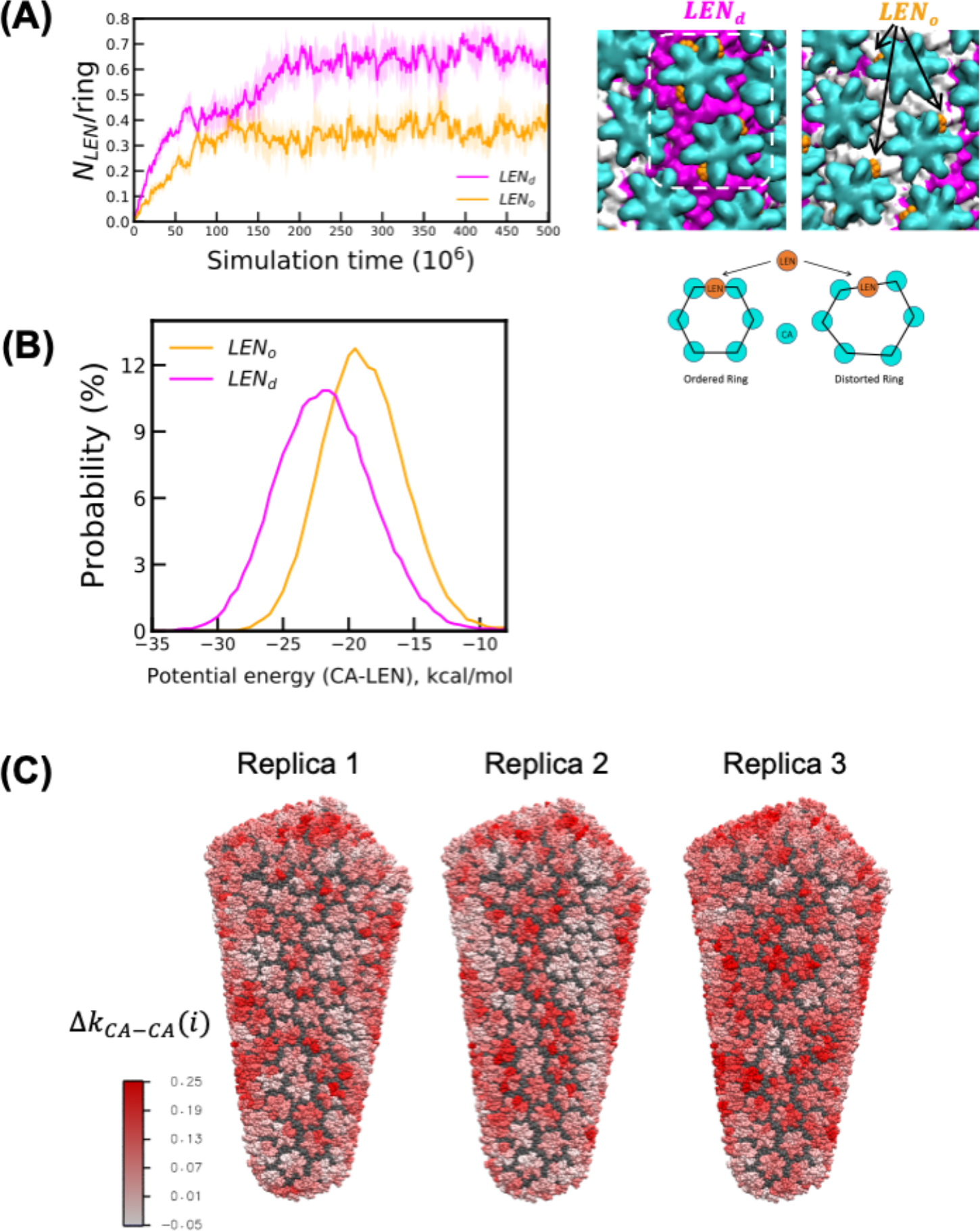
Molecular details of LEN binding and alteration of capsid microstructure. **(A)** The left panel shows the time series of the number of LEN molecules (normalized by the number of CA hexamer rings) bound to a CA monomer which is either part of the ordered lattice (*LEN*_*o*_) or distorted lattice (*LEN*_*d*_). The snapshot in the left-center panel shows LEN molecules bound to regions of the lattice that are ordered. The CTD domain of the CA monomers with ⟨*q*_6_⟩_*neigh*_ < 0.4 (distorted lattice) is colored in magenta. The CTD domain of the rest of the CA monomers (ordered lattice) is colored in white. The NTD domain of the CA monomer of all the hexamers is represented as cyan spheres. The highlighted region (orange box) shows 5 LEN molecules bound to 2 adjoining CA hexamers that are classified as distorted lattice. The snapshot in the right-center panel shows LEN bound to CA that are classified as ordered lattice. The right panel shows the schematic of LEN molecule (represented as orange circle) binding to ordered and distorted CA hexamer ring. **(B)** The probability distribution of the CA-LEN potential energy calculated for all LEN molecules bound to CA sites that are part of the ordered (*LEN*_*o*_) and distorted (*LEN*_*d*_) lattice. The CA-LEN potential energy calculations were performed for the final 250 × 10^6^ τ_*CG*_. **(C)** Deviation of the *k*_*CA*-*CA*_(*i*) for each CA monomer *i* relative to the value is calculated for the free capsid. The CA_NTD_ domain of all CA monomers for which the *k*_*CA*-*CA*_(*i*) increase and decrease relative to the free capsid is shown in red and blue, respectively. The red patches indicate effective stabilization relative to free capsid.

To determine if LEN binding modifies capsid stability, we represented the capsid as a heterogeneous elastic network model (hENM) (45). In the hENM model, the cumulative CA inter-subunit interactions are folded into an effective spring represented by harmonic potentials (Details in the *SI Appendix Methods*). The effective stability (represented as spring constants) is calculated using a rigorous method by first computing the normal modes of the elastic network model. In a way, the effective spring constants calculated from the normal mode analysis are the quantification of both flexibility and stability of the +LEN capsid CG structures. Note, the elasticity defined in this work is a molecular-level property, specifically focusing on the harmonic response of CA capsomers. By focusing on the harmonic response, we analyze how these capsomer building blocks in the lattice resist stress-induced dissociation. This determines if the capsid can squeeze through the narrow Nuclear Pore Complex (NPC) or if it will rupture.

A higher mean spring constant (*k*_*CA*-*CA*_) indicates stronger effective interactions. To establish a molecular view of the effective CA inter-subunit interactions, we calculated the deviation (Δ*k*_*CA*-*CA*_(*i*)) for CA subunit (*i*). The change in Δ*k*_*CA*-*CA*_(*i*) indicates the change in effective CA-CA interactions and illustrating how multiple CA hexamers are collectively stabilized (or destabilized) due to LEN binding (**Fig. 6C**). In the LEN-capsid complex, we observed the presence of spatially correlated CA subunits with positive Δ*k*_*CA*-*CA*_(*i*). The size of the hyperstabilized domains (**Fig. 6C**) ranged from 80-130 CA subunits, illustrating that LEN binding to the capsid induces hyperstabilized domains that are distributed across the hexameric lattice. Note that multiple LEN molecules are interspersed in these domains, cooperatively inducing hyperstabilization of the lattice.

## Discussion

In this study, we combined large-scale computer simulations, live-cell imaging, and structural analysis to characterize defect formation and rupture of LEN-treated HIV-1 cone-shaped capsids that are docking into the NPC central channel. We find that the substoichiometric binding of LEN to free capsids alters the properties of the capsid, specifically hyperstabilizing and rigidifying the lattice. CG MD simulations complemented by the outcome of live-cell imaging demonstrate that LEN-treated capsids dock at the NPC and rupture at the narrow end when bound to the central channel and then remain associated to the NPC after rupture. The dynamics of lattice rupture proceeds through three distinct steps: (1) disruption of CA-CA contacts at the hexamer-pentamer interface, (2) dissociation of CA pentamers at the narrow end, followed by at the wide end, and (3) nucleation of cracks at the hexamer-hexamer interface, typically in the vicinity of the narrow end. The viral cores are partly ruptured but not fully disassembled. We hypothesize that LEN-treated broken cores are stabilized by the interaction with the disordered FG-NUP98 mesh at the NPC. It is noteworthy that our recent transmission electron microscopy (analysis of HIV-1 cores isolated from virions revealed that treatment of the viral cores with LEN induced a high frequency of breaks at the narrow end of the capsids (42). These TEM results strongly support the CG MD simulations described in our studies – both docked at the NPC and free capsids.

The NPC allows selective and efficient transport of large cargoes between the cytoplasm and nucleus. The intact cone-shaped capsid is now understood to dock at the NPC central channel and translocate to the nuclear interior before disassembling (12-14). Our previous CG MD simulations of cone-shaped capsid docking into the NPC central channel demonstrated several key factors that regulate the passage of the intact capsid (16). When approaching from the narrow end, there is an increasing degree of steric barrier as the central channel encounters the wider ends of the capsid. One of the several behaviors through which the capsid lattice responds to steric stress is through the formation of striated patterns of greater lattice distortion (16, 35). Importantly, these patterns are signatures of the lattice elasticity and a necessary form of material metastability. This structural metastability is essential in maintaining capsid integrity during nuclear entry by absorbing radially inward stress from the NPC. It is to be noted that elasticity in this context refers to a CA monomeric-level effective harmonic response and not “macroscopic” elasticity. In other words, the elasticity in this work can also be interpreted as deviation of the ideal CA lattice from its ideal lattice structure is response to drug binding and confining stress. Our view is that this perspective is more useful for this problem than a macroscopic perspective as the capsid is, in fact, a mesoscopic object and not a macroscopic one. Structural analysis of LEN-treated capsids with substoichiometric concentrations of bound drug indicates the formation of hyperstabilized domains in the hexameric lattice inducing structural heterogeneity. As a consequence, the hexamer-pentamer interface is weakened relative to the hexamer-hexamer interface, resulting in the loss of pentamers, which is followed by nucleation of defects at the hexamer-hexamer interface. LEN binding stimulates similar changes in free capsids, but they occur with lower frequency on similar time scales. This suggests that the cores docked at the NPC are under increased stress, resulting in more frequent weakening of the hexamer-pentamer and hexamer-hexamer interactions, as well as more nucleation of defects at the hexamer-hexamer interface.

Once multiple pentamers are dissociated from the lattice, additional defects nucleate at hexamer-hexamer interfaces. Since all five pentamers are particularly important to maintain the high curvature of the narrow end, the loss of even a single pentamer can be detrimental to morphological integrity. The localized lattice rupture at the narrow end in our simulations with LEN is comparable with that of rupture induced by double-stranded reverse transcription products observed by atomic force microscopy (46-48). As the capsid breaks at the narrow end, defects at the hexamer-hexamer interface nucleate and extend from the narrow to the wide end. Importantly, these extended defects are formed along the striated patterns of lattice disorder, which we previously identified as weak points of the capsid (16, 35). At the endpoint of our CG MD simulations – and consistent with live-cell imaging – although there are extended cracks and vacancy-like defects in the lattice, suggesting loss of some of the capsid lattice, the capsid does not fully disassemble and retains a large part of the hexameric lattice. This observation is consistent with the visualization of damaged capsids with significant defects (49, 50). The partly broken hexameric lattice remains bound to the FG-NUP mesh at the NPC central channel. This suggests that the alteration in the morphology of the capsid lattice induced by LEN does not impair the capsid to FG-NUP interactions, allowing stable docking of the broken cores. These observations are also consistent with our observations of frequent breaks and damage induced by LEN treatment of isolated viral cores, as well as our observations in cell-based assays indicating that broken cores can dock at the NPC (42).

Multiple studies have reported two-step HIV-1 capsid uncoating: initial capsid rupture followed by terminal disassembly (13, 26, 51). The uncoating of broken capsids is delayed in the presence of host factors, which likely stabilizes the lattice (26, 27). Intrinsically disordered FG-rich NUP98 and NUP153 chains can oligomerize to higher-order condensates mediated by prion-like interactions and form a diffusion barrier for the capsid at the central channel and nuclear basket (30, 31, 52, 53). Whether the FG-NUP hydrogel can simultaneously stabilize the broken LEN-capsid complex at the NPC and prevent nuclear import is a direction for future investigation.

To summarize, our findings using large-scale modeling with supporting live-cell imaging reveal that the capsid inhibitor LEN modulates the HIV-1 capsid metastability and lattice molecular-scale properties. We find that FG-pocket binding of LEN cooperatively hyperstabilizes the hexameric lattice, inducing structural defects and then allowing breaking of the capsid, while altering capsid integrity overall. Lattice defects arising from the loss of pentamers and cracks along the weak points of the hexameric lattice drive the rupture of the capsid. Our results suggest that in the presence of the LEN, capsid docking into the NPC central channel will increase stress, resulting in more frequent breaks in the capsid lattice compared to free capsids. Lenacapavir modulates capsid structures by generating hyperstabilized domains, leading to detrimental loss of elasticity, essential for adaptation to the crowded environment at the NPC central channel during nuclear import. These insights into the effect of LEN on capsid lattice microstructure based on our modeling, and structural analysis can benefit future efforts in the rational design of more potent analogs.

## Material and Methods

Details of the CG models of NPC, HIV-1 capsid, RNP, and LEN are described in the *SI Appendix* (SI Methods S1-S3). Details of the CG MD simulations and analysis of these simulations are described in the *SI Appendix* (SI Methods S4-S5). Details of the system preparation of labeled viral cores for imaging, live-cell imaging experiments, and analysis of fluorescent virus particles are described in the SI Appendix (SI Methods S6).

## Data, Materials, and Software Availability

The study data are available upon request due to large file sizes and number of files.

## Acknowledgments

This research was supported in part by the National Institute of Allergy and Infectious Diseases (NIAID) of the National Institutes of Health (NIH) by grant U54 AI170855 for the Behavior of HIV in Viral Environments (B-HIVE) Center (to G.A.V.). This research was also supported in part by the Intramural Research Program of the NIH, National Cancer Institute, Center for Cancer Research (Z1A BC011436 to V.K.P.) and supplemental funding provided by Office of AIDS Research (to V.K.P.) to support collaborative interactions with the Behavior of HIV in Viral Environments Center (U54 AI170855)The coarse-grained simulations were performed using resources provided by the Frontera supercomputer at the Texas Advanced Computing Center (TACC) at the University of Texas at Austin and funded by the National Science Foundation (NSF) (grant OAC-1818253), as well as the Advanced Cyberinfrastructure Coordination Ecosystem: Services & Support (ACCESS) program, which is supported by NSF grant numbers 2138259, 2138286, 2138307, 2137603, and 2138296. Storage for simulation data was provided by TACC and the Research Computing Center (RCC) at the University of Chicago. The content of this publication does not necessarily reflect the views or policies of the Department of Health and Human Services, nor does mention of trade names, commercial products, or organizations imply endorsement by the US government.

## References

1. T. G. Müller, V. Zila, B. Müller, H.-G. Kräusslich, Nuclear Capsid Uncoating and Reverse Transcription of HIV-1. Annual Review of Virology 9, 261–284 (2022).

2. Q. Shen, C. Wu, C. Freniere, T. N. Tripler, Y. Xiong, Nuclear Import of HIV-1. Viruses 13 (2021).

3. B. K. Ganser, S. Li, V. Y. Klishko, J. T. Finch, W. I. Sundquist, Assembly and Analysis of Conical Models for the HIV-1 Core. Science 283, 80–83 (1999).

4. S. Li, C. P. Hill, W. I. Sundquist, J. T. Finch, Image reconstructions of helical assemblies of the HIV-1 CA protein. Nature 407, 409–413 (2000).

5. J. A. G. Briggs et al., The stoichiometry of Gag protein in HIV-1. Nature Structural & Molecular Biology 11, 672–675 (2004).

6. M. Gupta, A. J. Pak, G. A. Voth, Critical mechanistic features of HIV-1 viral capsid assembly. Science Advances 9, eadd7434 (2023).

7. S. Mosalaganti et al., AI-based structure prediction empowers integrative structural analysis of human nuclear pores. Science 376, eabm9506 (2022).

8. A. E. Hulme, O. Perez, T. J. Hope, Complementary assays reveal a relationship between HIV-1 uncoating and reverse transcription. Proceedings of the National Academy of Sciences 108, 9975–9980 (2011).

9. J. I. Mamede, G. C. Cianci, M. R. Anderson, T. J. Hope, Early cytoplasmic uncoating is associated with infectivity of HIV-1. Proceedings of the National Academy of Sciences 114, E7169–E7178 (2017).

10. R. C. Burdick et al., Dynamics and regulation of nuclear import and nuclear movements of HIV-1 complexes. PLOS Pathogens 13, e1006570 (2017).

11. A. Dharan, N. Bachmann, S. Talley, V. Zwikelmaier, E. M. Campbell, Nuclear pore blockade reveals that HIV-1 completes reverse transcription and uncoating in the nucleus. Nature Microbiology 5, 1088–1095 (2020).

12. R. C. Burdick et al., HIV-1 uncoats in the nucleus near sites of integration. Proceedings of the National Academy of Sciences 117, 5486–5493 (2020).

13. C. Li, R. C. Burdick, K. Nagashima, W.-S. Hu, V. K. Pathak, HIV-1 cores retain their integrity until minutes before uncoating in the nucleus. Proceedings of the National Academy of Sciences 118, e2019467118 (2021).

14. V. Zila et al., Cone-shaped HIV-1 capsids are transported through intact nuclear pores. Cell 184, 1032-1046.e1018 (2021).

15. K. Jan Philipp et al., Passage of the HIV capsid cracks the nuclear pore. bioRxiv 10.1101/2024.04.23.590733, 2024.2004.2023.590733 (2024).

16. A. Hudait, G. A. Voth, HIV-1 capsid shape, orientation, and entropic elasticity regulate translocation into the nuclear pore complex. Proceedings of the National Academy of Sciences 121, e2313737121 (2024).

17. S. M. Bester et al., Structural and mechanistic bases for a potent HIV-1 capsid inhibitor. Science 370, 360–364 (2020).

18. J. O. Link et al., Clinical targeting of HIV capsid protein with a long-acting small molecule. Nature 584, 614–618 (2020).

19. P. C. Patel, H. K. Beasley, A. Hinton, C. N. Wanjalla, Lenacapavir (Sunlenca®) for the treatment of HIV-1. Trends in Pharmacological Sciences 44, 553–554 (2023).

20. D. A. Bejarano et al., HIV-1 nuclear import in macrophages is regulated by CPSF6-capsid interactions at the nuclear pore complex. eLife 8, e41800 (2019).

21. A. C. Francis et al., HIV-1 replication complexes accumulate in nuclear speckles and integrate into speckle-associated genomic domains. Nature Communications 11, 3505 (2020).

22. G. A. Sowd et al., A critical role for alternative polyadenylation factor CPSF6 in targeting HIV-1 integration to transcriptionally active chromatin. Proceedings of the National Academy of Sciences 113, E1054–E1063 (2016).

23. Christopher R. Chin et al., Direct Visualization of HIV-1 Replication Intermediates Shows that Capsid and CPSF6 Modulate HIV-1 Intra-nuclear Invasion and Integration. Cell Reports 13, 1717–1731 (2015).

24. G. Wei et al., Prion-like low complexity regions enable avid virus-host interactions during HIV-1 infection. Nature Communications 13, 5879 (2022).

25. J. C. V. Stacey et al., Two structural switches in HIV-1 capsid regulate capsid curvature and host factor binding. Proceedings of the National Academy of Sciences 120, e2220557120 (2023).

26. K. M. R. Faysal et al., Pharmacologic hyperstabilisation of the HIV-1 capsid lattice induces capsid failure. eLife 13, e83605 (2024).

27. R. C. Burdick et al., HIV-1 uncoating requires long double-stranded reverse transcription products. Science Advances 10, eadn7033 (2024).

28. S. Petrovic et al., Architecture of the linker-scaffold in the nuclear pore. Science 376, eabm9798 (2022).

29. S. C. Ng et al., Barrier properties of Nup98 FG phases ruled by FG motif identity and inter-FG spacer length. Nature Communications 14, 747 (2023).

30. L. Fu et al., HIV-1 capsids enter the FG phase of nuclear pores like a transport receptor. Nature 626, 843–851 (2024).

31. C. F. Dickson et al., The HIV capsid mimics karyopherin engagement of FG-nucleoporins. Nature 626, 836–842 (2024).

32. J. Jin, A. J. Pak, A. E. P. Durumeric, T. D. Loose, G. A. Voth, Bottom-up Coarse-Graining: Principles and Perspectives. Journal of Chemical Theory and Computation 18, 5759–5791 (2022).

33. J. P. Kreysing et al., Passage of the HIV capsid cracks the nuclear pore. Cell 188, 930-943.e921 (2025).

34. A. Deshpande et al., Elasticity of the HIV-1 core facilitates nuclear entry and infection. PLoS Pathog 20, e1012537 (2024).

35. A. Yu et al., Strain and rupture of HIV-1 capsids during uncoating. Proceedings of the National Academy of Sciences 119, e2117781119 (2022).

36. Z. Hou et al., HIV-1 nuclear import is selective and depends on both capsid elasticity and nuclear pore adaptability. Nature Microbiology 10, 1868–1885 (2025).

37. M. Yu et al., Visualizing the disordered nuclear transport machinery in situ. Nature 617, 162–169 (2023).

38. E. E. Najbauer, S. C. Ng, C. Griesinger, D. Görlich, L. B. Andreas, Atomic resolution dynamics of cohesive interactions in phase-separated Nup98 FG domains. Nature Communications 13, 1494 (2022).

39. C. M. Highland, A. Tan, C. L. Ricaña, J. A. G. Briggs, R. A. Dick, Structural insights into HIV-1 polyanion-dependent capsid lattice formation revealed by single particle cryo-EM. Proceedings of the National Academy of Sciences 120, e2220545120 (2023).

40. D. Singh et al., The molecular architecture of the nuclear basket. Cell 187, 5267-5281.e5213 (2024).

41. A. Barducci, G. Bussi, M. Parrinello, Well-Tempered Metadynamics: A Smoothly Converging and Tunable Free-Energy Method. Physical Review Letters 100, 020603 (2008).

42. C. Li et al., Lenacapavir disrupts HIV-1 core integrity while stabilizing the capsid lattice. Proceedings of the National Academy of Sciences 122, e2420497122 (2025).

43. W. Lechner, C. Dellago, Accurate determination of crystal structures based on averaged local bond order parameters. The Journal of Chemical Physics 129, 114707 (2008).

44. P. J. Steinhardt, D. R. Nelson, M. Ronchetti, Bond-orientational order in liquids and glasses. Physical Review B 28, 784–805 (1983).

45. E. Lyman, J. Pfaendtner, G. A. Voth, Systematic Multiscale Parameterization of Heterogeneous Elastic Network Models of Proteins. Biophys. J. 95, 4183–4192 (2008).

46. S. Rankovic, J. Varadarajan, R. Ramalho, C. Aiken, I. Rousso, Reverse Transcription Mechanically Initiates HIV-1 Capsid Disassembly. Journal of Virology 91, 10.1128/jvi.00289-00217 (2017).

47. S. Rankovic, R. Ramalho, C. Aiken, I. Rousso, PF74 Reinforces the HIV-1 Capsid To Impair Reverse Transcription-Induced Uncoating. Journal of Virology 92, 10.1128/jvi.00845-00818 (2018).

48. S. Rankovic, A. Deshpande, S. Harel, C. Aiken, I. Rousso, HIV-1 Uncoating Occurs via a Series of Rapid Biomechanical Changes in the Core Related to Individual Stages of Reverse Transcription. Journal of Virology 95, 10.1128/jvi.00166-00121 (2021).

49. A. Guedán et al., HIV-1 requires capsid remodelling at the nuclear pore for nuclear entry and integration. PLOS Pathogens 17, e1009484 (2021).

50. A. C. Francis et al., Localization and functions of native and eGFP-tagged capsid proteins in HIV-1 particles. PLOS Pathogens 18, e1010754 (2022).

51. C.L. Márquez et al., Kinetics of HIV-1 capsid uncoating revealed by single-molecule analysis. eLife 7, e34772 (2018).

52. S. Milles, Edward A. Lemke, Single Molecule Study of the Intrinsically Disordered FG-Repeat Nucleoporin 153. Biophysical Journal 101, 1710–1719 (2011).

53. S. Milles et al., Facilitated aggregation of FG nucleoporins under molecular crowding conditions. EMBO reports 14, 178–183 (2013).

