## Supporting Information for "Lenacapavir-induced Lattice Hyperstabilization is Central to HIV-1 Capsid Failure at the Nuclear Pore Complex and in the Cytoplasm"

**This file contains SI Materials and Methods, SI Figures 1-8, SI Movie Legends 1-5**

### SI Methods

#### S1. CG model of NPC.

To overcome the computational cost associated with simulating the Nuclear Pore Complex in fully atomistic detail, we recently developed a coarse-grained (CG) model from available experimental structural data (1). The model is largely "bottom-up" i.e., developed quantitatively from atomistic simulations of constituent nucleoporin subcomplexes. We used a mapping resolution of ~ 5 amino acids per CG site or "bead". The composite CG model of the NPC consists of cytoplasmic ring (CR), nuclear ring (IR), and inner ring (IR). The CR and NR consist of 8 copies of Y-complex dimer. IR consists of 8 subunits, and each subunit is composed of 2 copies each of NUP155, NUP93, NUP205, and NUP188. The linker NUP35 connects adjacent subunits. 2 additional copies of NUP155 also connect each IR subunit to NUP160 of Y-complex of CR and NR. Our CG model also consists of the capsid-binding coiled-coil NUP62, and intrinsically disordered NUP98 chains. These proteins contain phenylalanine-glycine (FG) motifs that bind to a hydrophobic pocket in capsid (CA) monomer. NUP62 is modeled as a heterotrimeric NUP54-NUP58-NUP62 subcomplex connected using a weakly bonded harmonic network with a force constant of 0.01 kcal mol$^{-1}$Å$^{-2}$ between neighboring monomers with a distance cutoff of 3 nm.

Before deriving the inter and intra-molecular interactions of the CG proteins, each constituent nucleoporin (NUP) protein of the NPC was mapped to corresponding CG beads using the essential dynamics coarse-graining (EDCG) method (2), in which the atomistic residues are grouped to CG beads such that the principal modes of motions sampled during atomistic simulations are preserved. In EDCG, the mapping operator ($\boldsymbol{M}_{\mathbf{R}}^{N}: \mathbf{r}^{n} \rightarrow \mathbf{R}^{N}$), is variationally optimized using simulated annealing to obtain the global minimum of the target residual ($\chi^2$). Here, $\mathbf{r}^{n}$ and $\mathbf{R}^{N}$ are the configurations of the atomistic and coarse-grained trajectories, respectively. The all-atom to coarse-grained mapping function is adjusted during the optimization to minimize the target residual ($\chi^2$):

$$\chi^2 = \frac{1}{3N} \sum_{I=1}^{N} \langle \sum_{i,j} \left| \boldsymbol{r}_i - \boldsymbol{r}_j \right|^2 \rangle_t \quad : \ i,j \in I, j \geq i \qquad \text{(S1)}$$

where, $N$ is the total number of CG sites. Here $i,j$ are the unique pairs in the group of all-atom residues that are part of the CG site, $I$. $\boldsymbol{r}_i = \boldsymbol{x_i} - \langle\, \boldsymbol{x_i} \,\rangle_t$ is the displacement of atom $i$ from the atom's mean position, $\langle\, \boldsymbol{x_i} \,\rangle_t$. The residual ($\chi^2$) is small when the atoms $i, j$ move in a correlated fashion, i.e. the displacements $\boldsymbol{r}_i$ and $\boldsymbol{r}_j$ are similar.

By design, it is easier to minimize the residual ($\chi^2$), if a single residue in atomistic resolution is mapped to one CG bead (i.e. 1→1 mapping), and reverse when very coarse mapping is used. However, computational cost also increases for finer coarse-grained models. Therefore, our choice of the mapping resolution (i.e., average number of residues per CG beads) is based on an empirical "elbow rule", i.e., the $\chi^2$ value with increasing value of $N$, the $\chi^2$ value decreases and reaches a plateau. We chose the $N$ value at the elbow point, at which $\chi^2$ reaches the plateau.

The intermolecular non-bonded CG interactions between NUP monomers and subcomplexes were modeled using repulsive excluded volume and attractive interactions. The repulsive excluded volume interactions were used to prevent overlap between CG beads. The repulsive excluded volume interaction ($E_{excl}$) was modeled with a soft cosine potential,

$$E_{excl}(r_{ij}) = A\left(1 + \cos\left(\frac{\pi r_{ij}}{r_c}\right)\right) \tag{S2}$$

where $r_{ij}$ is the pairwise distance between CG site types $i$ and $j$. $A$ is 15 kcal/mol for all $ij$ pairs. The distance cutoff ($r_c$) for the excluded volume interactions was 1.0 nm. The non-bonded attractive interactions between NUP subcomplexes at the key binding interactions were modeled with pairwise Gaussian potential ($E_{gauss}$),

$$E_{gauss}(r_{ij}) = \frac{H_{ij}}{\sigma_{ij}\sqrt{2\pi}} \exp\left(-\frac{\left(r_{ij} - r_{0,ij}\right)^2}{2\sigma_{ij}^2}\right) \tag{S3}$$

where $r_{0,ij}$ and $\sigma_{ij}$ are the mean and standard deviation of the distance between CG site types $i$ and $j$. For all CG $ij$ pairs $\sigma_{ij}$ value of 0.12 nm was used for all non-bonded attractive interactions modeled with $E_{gauss}$. The distance cutoff of 3 nm is used for all The constant $H_{ij}$ for each binding interface was optimized using Relative Entropy Minimization from the atomistic trajectories of the respective all-atom subcomplexes (3).

CG NUP98$_{1\text{-}620}$ chains consist of the structured GLEB domain (residue 157-213) and the intrinsically disordered region (residue 1-156 and residue 214-620) modeled as a linear polymer chain. Each NUP98$_{1\text{-}620}$ consists of 124 CG beads. Each CG bead of the polymer chain is linked with a harmonic spring with equilibrium bond length of 2 nm and force constant of 0.5 kcal/mol Å$^{-2}$. The intrinsically disordered regions (residue: 1-156 and residue: 214-480) predominantly consist of the capsid-binding FG-motif. The intra-chain and inter-chain interactions between CG beads of the FG-rich region (residue: 1-156 and residue: 214-480) were modeled using a 12-6 Lennard-Jones potential ($E_{scLJ}$) with a modified soft-core (4):

$$E_{scLJ}(r) = 4\varepsilon\lambda^n \left[1/\left(\alpha_{LJ}(1-\lambda)^2 + (r/\sigma)^6\right)^2 - 1/\left(\alpha_{LJ}(1-\lambda)^2 + (r/\sigma)^6\right)\right] \tag{S4}$$

where, $n$ = 2, $\alpha_{LJ}$ = 0.5, $\alpha$ = 0.6, and $\sigma$ = 1.25 nm. The distance cutoff of 3 nm was used. The strength ($\epsilon$) of the inter- and intra-chain interactions (also denoted as $\varepsilon_{NUP98}$ in the main manuscript) is used as 0.3 kcal/mol. A total of 48 NUP98 chains tethered to NUP155 proteins at the IR.

The protocol to create and equilibrate the composite NPC-membrane system is described in our previous publication (5). The dimension of the lipid bilayer in the $x$ and $y$ direction is 200 nm. The CG lipid was modeled using a 4-site model consisting of 1 head, 1 interfacial, and 2 hydrophobic tail beads using the same potential energy function as in ref. (6). Membrane-binding interactions of NPC are modeled with $E_{scLJ}$. Here, $n$ = 2, $\alpha_{LJ}$ = 0.5, $\alpha$ = 0.6, and $\sigma$ = 1.5 nm. The distance cutoff of 4 nm was used. Protein-lipid interactions were added between the CG sites of the $\beta$-propeller domain of NUP155, NUP133, and NUP160, and the CG lipid head group.

**S2. CG model of HIV-1 Capsid and RNP.**

The CG model of HIV-1 capsid was derived from an atomistic simulation trajectory of a composite system of three CA hexamers complexed with IP6 and FG peptides. The details of the atomistic simulation and CG model development protocol are described in ref. (5). Each CA monomer consist of 46 CG sites with a mapping resolution of ~5 AA residues per CG site. The intramonomer bonding topology of the CA monomer is modeled as a heterogeneous elastic network with a radial cutoff of 3 nm. The non-bonded interactions between CA monomers are modeled with a combination of repulsive excluded volume ($E_{excl}$) and attractive pairwise Gaussian interactions ($E_{gauss}$). The value of $A$ in $E_{excl}$ was 15 kcal/mol for all $ij$ pairs of CA. The distance cutoff ($r_c$) for the excluded volume interactions was set at 1.0 nm. The parameters $r_{0,ij}$ and $\sigma_{ij}$ of $E_{gauss}$ were determined from the atomistic simulation trajectory. The constant $H_{ij}$ for all CG $ij$ pairs was optimized using Relative Entropy Minimization from the corresponding atomistic simulation trajectory (3). The parameters of $E_{gauss}$ for the non-bonded attractive interactions between CA and the FG peptide were derived using the identical protocol as CA-CA interactions.

The ribonucleoprotein complex (RNP) within the capsid is modeled as a linear polymer chain containing 3000 beads. In our simulations there are RNP chains. The non-bonded interactions between CA monomers are modeled with $E_{scLJ}$. Here, $n$ = 2, $\alpha_{LJ}$ = 0.5, $\alpha$ = 0.6, and $\sigma$ = 1.25 nm. The condensation state of RNP was regulated by varying the strength of the RNP-RNP interaction ($\varepsilon_{RNP}$). To model uncondensed RNP and condensed RNP, $\varepsilon_{RNP}$ values of 0.3 kcal/mol and 0.7 kcal/mol were used. To emulate electrostatic interactions between RNP and the basic residues at the C-terminal of CA, we used attractive pairwise Gaussian interactions ($E_{gauss}$) between the RNP and charged residues in the C-terminal end of CA ($H_{ij}$ = - 4.5 kcal/mol, $r_{0,ij}$ = 1.25 nm, and $\sigma_{ij}$ = 0.12 nm).

**S3. CG model of Lenacapavir.**

The CG model of LEN and LEN-CA molecular interactions was derived directly from the experimental X-ray crystal structure of 6 LEN molecules complexed to a CA hexamer (PDB: 6VKV) (7). The CG LEN model consists of 8 sites: 1. $CG_1$ – pyridinium ring (R1), 2. $CG_2$ – 2-(methanesulfonyl)-2-methylpropane group attached to the pyridinium ring (R1), 3. $CG_3$ – indazole ring (R2), 4. $CG_4$ – trifluoroethyl ($-CH_2-CF_3$) group attached to the indazole ring (R2), 5. $CG_5$ – sulfonamide group attached to the indazole ring (R2), 6. $CG_6$ – difluorobenzyl ring (R3), 7. $CG_7$ – acetamide group attached to the pyridinium ring (R1), 8. $CG_8$ – cyclo-pentapyrazole ring (R4). The CG Lenacapavir is modeled as a rigid body, i.e., without any intramolecular fluctuations. Lenacapavir interacts with two adjoining CA monomers at the binding pocket through hydrogen bonding and Van der Waals interactions (7). Specifically, all the ring systems of Lenacapavir interact with multiple residues of $CA_1$-NTD and $CA_2$-CTD through extensive Van der Waals interactions. Additionally, Lenacapavir forms hydrogen bonding interactions with N57, K70, and N74 of $CA_1$-NTD, S41 of $CA_2$-NTD, and Q179 and N183 of $CA_2$-CTD. Specifically, the sulfonyl group of the R1 ring and the sulfonamide group of the R2 ring form the hydrogen-bonding network of interactions with the CA subunits.

Note, our decision to derive the non-bonded interactions of the LEN CG model directly from the experimental crystal structure is due to the lack of a reliable atomistic force field. Specifically, small changes in the force field parameters of the electrostatic groups of Lenacapavir lead to unrealistic variation of the affinity or dissociation free energy of the drug molecule to CA. is due to the lack of a reliable atomistic force field. Lack of a reliable atomistic force field is also a barrier in deriving reliable intramolecular CG topology (bonds, angles, and harmonic force constants).

The CG interactions between CA and LEN were modeled using pairwise repulsive excluded volume ($E_{Excl}$) and attractive ($E_{Gauss}$) interactions. The distance cutoff ($r_c$) for the excluded volume interactions between CA-LEN and LEN-LEN interactions were set at 1.25 nm. The value of $A$ was set to 15 kcal/mol for all $ij$ pairs. In our CG model, the LEN-CA associative interactions are modeled using two different energy scales to model the hydrogen bonding ($H_{ij,HB}$) and Van der Waals interactions ($H_{ij,VDW}$). First, we mapped the 6 LEN molecules complexed to a CA hexamer (PDB: 6VKV) from atomistic to coarse-grained resolution. We first assigned $H_{ij,VDW}$ between LEN CG beads and CA CG beads to mimic the extensive Van der Waals interactions between LEN ring systems and CA residues. Then we assigned $H_{ij,HB}$ between LEN CG beads and CA CG beads to mimic the hydrogen bonded interactions between LEN sulfonyl (R1) and sulfonamide (R2) groups and CA residues. The $r_{0,ij}$ values of the LEN-CA CG interactions were derived directly from the CG-mapped structure of 6 LEN molecules complexed to CA hexamer. In the simulations, $H_{ij,VDW}$ is chosen (-6.25 kcal mol$^{-1}$ nm$^{-1}$) such that when $H_{ij,HB}$ is turned off, multiple LEN molecules continuously associate and dissociate at the CA hexamer binding pocket. In other words, in the CG LEN model, the energy scale of the Van der Waals interactions is not strong enough to initiate the association of LEN to the capsid. The value of $H_{ij,HB}$ (-8.50 kcal mol$^{-1}$ nm$^{-1}$) is the minimum CA-LEN binding strength required to initiate LEN association. Finally, in our simulations, all attractive interactions between LEN and CA monomers that are part of the pentamer ring are turned off, as in experiments, LEN selectively associates to CA hexamers (8).

**S4. Coarse-grained Simulations.**

All CG MD simulations were performed using the large-scale atomic/molecular massively parallel simulator (LAMMPS) (9). The details of the CG models of different components (NPC, lipid, HIV-1 capsid, RNP, and LEN) are described in the *SI Appendix* and in ref. (5). The details of the construction of the composite membrane-embedded NPC are provided in ref. (5). The CG MD simulations were performed using time step ($\tau_{CG}$) 50 fs. The equations of motion were integrated with the Velocity Verlet algorithm, and the simulation setup was periodic in all $(x, y, z)$ dimensions. In the initial system, the cone-shaped LEN-capsid complex was placed such that the narrow end is coplanar to the cytoplasmic Y-complex, and the axis of the cone is the same as the $z$ axis. Note the $(x, y)$ plane is the plane of the nuclear membrane. An additional 800 LEN molecules are randomly placed in the void region of the simulation cell, specifically above the nuclear membrane, both at the cytoplasmic and nuclear sides. The production simulations of NPC central channel were performed in the constant $Np_{xy}T$ ensemble at 310 K and 1 bar. The temperature of the system was maintained using Langevin thermostat with a coupling constant 2000 $\tau_{CG}$ (10), and pressure was maintained using a Nose-Hoover barostat with a coupling constant of 4000 $\tau_{CG}$ (11). In the CG simulations, LEN is modeled as a rigid body. Therefore, the translational and rotational motions of LEN molecules are modeled using rigid body dynamics (12, 13).

The non-NPC simulations of LEN binding to the capsid were performed in a simulation cell of dimension. 300 nm in $x$, $y$, and $z$ directions. The periodic boundary condition is used in all directions. Initially, the capsid is positioned such that the geometric center of the capsid is the same as the center of the simulation box. 600 LEN molecules were initially randomly positioned in the void space of the simulation cell, such that LEN molecules are at least 2 nm away from the capsid surface. The simulations were performed at 310 K, and the temperature was maintained using Langevin thermostat with a coupling constant of 1000$\tau_{CG}$ (10). Simulation trajectory snapshots were saved every 50000 $\tau_{CG}$.

We performed WTMetaD simulations to facilitate the rupture of HIV-1 in non-NPC simulations (14). These simulations mimic *in vitro* conditions. To define the collective variable (CV) for WTMetaD simulation, we calculated the number of CA monomers in hexamer rings ($CA_{hex}$) that are direct neighbors of the CA monomers in pentamer rings ($CA_{pen}$). To this end, we calculate the geometric center of the CG beads of the C-terminal domain. The number of adjacent $CA_{hex}$ for each $CA_{pen}$ was calculated with a radial cutoff of 4.0 nm, considering the geometric center of the CTD of each CA monomer. Then, we calculate the mean value of $CA_{pen}$ and apply bias to this collective variable ($CN_{Pen-Hex}$). The mean coordination number function decays to zero between 4.0 nm and 4.5 nm. The time-series of the CV values are shown in Fig. S6. For computational efficiency, we set the upper limit of CV, a weak harmonic restraint of force constant 10000 kcal/mol/nm$^2$ to prevent the CV value from exceeding 2.5. In other words, this harmonic restraint is necessary to prevent over-coordination of CTD domains. The height of Gaussian biases was set to 0.6 kcal/mol and was deposited every $500\tau_{CG}$ with a bias factor of 100 and width of 0.1.

#### S5. Analysis of CG MD Simulations.

The trajectories of the CG MD simulations were visualized, and the simulation snapshots were prepared in Visual Molecular Dynamics (VMD) software (15).

(a) CA-LEN binding analysis: To determine if a LEN molecule is bound to CA monomers in the simulations, we used the following criteria:

$$d_{CA-LEN} = \left|\frac{1}{N_{CA}}\sum_{i=1}^{N} r_{CA(i)} - \frac{1}{N_{LEN}}\sum_{i=1}^{N} r_{LEN(i)}\right| \quad \text{(S5)}$$

where $d_{CA-LEN}$ is the distance between the center of mass of the selected CG sites of LEN and CA. $r$ is the coordinate of the selected CG sites of CA and LEN. For the calculation of the center of mass of LEN, the CG beads 1, 3, 6, and 8 are used, which correspond to the ring systems R1, R2, R3, and R4. For the calculation of the center of mass of CA, the CG beads 8 (residue: 34-40), 10-13 (residue: 48-76), 21-22 (all-atom residue: 99-105), 34-37 (residue: 170-182) were used. A CA monomer and LEN molecule are in contact if $d_{CA-LEN} < 3\ nm$. To determine $N_{bound}$ (number of LEN molecules bound to CA hexamers), we only consider LEN molecules that simultaneously contact 2 adjoining CA monomers of a hexamer.

(b) Capsid lattice order analysis: To determine the lattice order of the capsid, specifically whether constituent CA monomers of a ring are geometrically ideal or distorted, we used the neighbor averaged ($\langle q_6 \rangle_{neigh}$) Steinhardt bond order parameter (16, 17). First, we created a reduced form of the CG MD simulation trajectories by selecting the CG site 21 (residues: 99-105) of each CA monomer of the capsid. We note that CG site 21 is approximately the geometric center of a CA monomer. Therefore, in the reduced form, the capsid consists of 1314 beads which is the same as the number of CA monomers. Using the reduced form of the trajectory, the neighbor averaged Steinhardt bond order parameter of a CA monomer is calculated as follows:

$$\langle q_6 \rangle_{neigh} = \left(\frac{4\pi}{2l+1}\sum_{m=-l}^{l}|\bar{q}_{lm}|^2\right)^{1/2} \quad \text{(S6)}$$

where $\bar{q}_{lm}(i) = \frac{1}{N_B(i)}\sum_{k=0}^{N_B(i)} q_{lm}(i)$ and $l = 6$. Here $N_B$ is the number of neighbors of CA monomer $i$, and the reference CA monomer itself. The term $q_{lm} = \frac{1}{N_B}\sum_{i=1}^{N_B} \Upsilon_{lm}(\theta_i(r), \phi_i(r))$, and $\Upsilon_{lm}(\theta_i(r), \phi_i(r))$ are spherical harmonics of rank *l* and *m*, where $\theta_i(r)$ and $\phi_i(r)$ are the polar angles of each of the $N_B$ bonds between the central CA monomer and neighboring CA monomers. From a central CA monomer $i$ we consider the three closest CA neighbors. We defined a CA monomer to be in a geometrically ordered environment if $\langle q_6 \rangle_{neigh} > 0.4$.

(c) Capsid elasticity analysis: To evaluate how LEN binding modulates capsid stiffness, we used the heterogeneous elastic network model (hENM) method (18). In the hENM framework, harmonic bonds are assigned during the analysis to all CG site pairs $ij$ within a cutoff distance from the central CG site $i$. The harmonic force constants ($k_{ij}$) for each bond have the same value, which are then iteratively optimized by computing the normal modes of the elastic network model, solving the eigenvalue problem,

$$\boldsymbol{H}\boldsymbol{v}_{\boldsymbol{k}} = \omega_k^2 \boldsymbol{M}\boldsymbol{v}_{\boldsymbol{k}} \tag{S7}$$

where, $H$ is the Hessian: $H_{i,j} = \left.\frac{\partial^2 V}{\partial q_i \partial q_j}\right|_m$, $\boldsymbol{M}$ is the diagonal matrix for the masses of the particles, and $\omega_k$ frequency for the mode of motion. The solution to the equation of motion

$$\boldsymbol{M}\frac{\mathrm{d}^2\boldsymbol{q}}{\mathrm{d}t^2} + \boldsymbol{H}\boldsymbol{q} = 0 \tag{S8}$$

where $\boldsymbol{q}$ is the generalized coordinate, and that for $N$ classically, interacting particles near the potential energy minimum, $\boldsymbol{q}_m$:

$$V(\boldsymbol{q}) = V(\boldsymbol{q}_m) + \sum_i \left.\frac{\partial V}{\partial q_i}\right|_m (q_i - q_{i,m}) + \frac{1}{2}\sum_{i,j} \left.\frac{\partial^2 V}{\partial q_i \partial q_j}\right|_m (q_i - q_{i,m})(q_j - q_{j,m}) + O(\boldsymbol{q} - \boldsymbol{q}_m)^3 \tag{S9}$$

$V(\boldsymbol{q}_m)$ is a constant, and $\left.\frac{\partial V}{\partial q_i}\right|_m$ is zero. Using the normal modes, the amplitudes are then scaled according to equipartition energy that reflects the temperature of the reference simulations. The harmonic force constants ($k_{ij}$) are then iteratively optimized so that mean-squared fluctuations of the coarsened system ($\langle r_{ij}^2 \rangle = \langle (x_{ij} - \langle x_{ij} \rangle)^2 \rangle$) match that of the fine-grained reference system

$$\frac{1}{k_{ij}^{n+1}} = \frac{1}{k_{ij}^{n}} - \alpha\left(\langle r_{ij}^2 \rangle_{\mathrm{CG}} - \langle r_{ij}^2 \rangle_{\mathrm{ref}}\right) \tag{S10}$$

Here, $n$ is the number of iterations and $\alpha$ is a parameter that controls the scale of the adjustment in the spring constant for each iteration.

First, from the LEN binding simulations to the capsid, we picked configurations of LEN complexed to the capsid with monotonically increasing $N_{bound}$. The unbound LEN molecules were removed from the simulation box. The LEN bound to capsid complexes at different values of $N_{bound}$ was then further evolved for 25 × $10^6$ $\tau_{CG}$. Then, we generated a reduced version of the simulation trajectory by selecting CG site 21 (geometric center) of each CA monomer. Then, the averaged trajectory was computed by aligning all the trajectory snapshots to the initial configuration. We choose a distance cutoff of 5 nm, which typically connects the central CG site with the closest 3 neighbors (3 $ij$ pairs). The hENM analysis was performed from the final 20 × $10^6$ $\tau_{CG}$ of the reduced version of the trajectory. The capsid stiffness parameter ($k_{CA-CA}$) is then calculated as the mean harmonic force constant for all $ij$ bonds between CA monomers.

**S6. Live-cell imaging assay.**

HeLa:POM121-HALO cells and human embryonic kidney 293T cells (ATCC CRL-3216) were maintained in Dulbecco's modified Eagle's medium (DMEM) supplemented with 10% fetal calf serum (Hyclone) and 1% penicillin-streptomycin (50 units and 50 µg/ml, respectively; hereafter referred to as Complete Media). All cells were maintained in a humidified incubator at 37°C with 5% $CO_2$. HeLa:POM121-HALO cells were generated by transduction of HeLa cells with VSV-G-pseudotyped virions containing a lentiviral vector expressing the C-terminal HALO-tagged human POM121. This lentiviral construct was generated by replacing mCherry with HALO in a lentiviral vector that expresses human POM121-mCherry under the control of the ubiquitin C promoter (19).

Virions labeled with GFP-CA and cmHALO were prepared by polyethylenimine transfection (PEI; Polysciences) of 293T cells (3.5 x $10^6$ cells seeded in 100-mm cell culture dish one day prior) with the HIV-1-based vectors pHmNG (40% of the total HIV-1 plasmid amount), pHGFP-GFPCA (10% of the total HIV-1 plasmid amount), and pHmNG-iHALO (50% of the total HIV-1 plasmid amount), along with pHCMV-G (20), which expresses the G glycoprotein of vesicular stomatitis virus (VSV-G). pHmNG, which contains an mNeonGreen (mNG) reporter gene in place of *nef* and does not express *env*, was generated by replacing the GFP reporter gene in pHGFP (21) with mNG. pHGFP-GFPCA (22) is similar to pHmNG, except that a GFP-CA fusion protein is generated after proteolytic processing of Gag and there is a GFP reporter gene in place of *nef*. pHmNG-iHALO expresses free HALO and CA after proteolytic processing and there is an mNG reporter gene in place of *nef*. To construct pHmNG-iHALO, the GFP in HIV Gag-iGFP ΔEnv (NIH AIDS Reagent Program, Division of AIDS, NIAID, NIH, from Dr. Benjamin Chen; Cat#12455) was replaced with HALO and then a portion of gag containing HALO was transferred to pHmNG. Three hours after transfection, Complete Media containing 20 nM Janelia Farm 646 (JF646) dye was added to the cells. Unlabeled virions were prepared by PEI transfection of 293T cells with pHmNG and pHCMV-G. Supernatants from the transfected cells were collected 24 hours after transfection, clarified using a 0.45 μm membrane filter, and concentrated by ultracentrifugation (100,000 × g) for 1.5 hours at 4°C through a 20% sucrose cushion (wt/vol) in 1X Dulbecco's Phosphate-Buffered Saline with calcium and magnesium (PBS). The concentrated virus was resuspended in 500 μl Complete Media or PBS and the virus amounts were determined by p24 ELISA (XpressBio).

Viral lysates were prepared using Pierce RIPA buffer (ThermoFisher) and subjected to western blot analysis using mouse anti-p24 antibody (NIH Aids Reagent Program Cat#24-3), rabbit anti-GFP (ThermoFisher Cat#A6455), and rabbit anti-HALO (Promega Cat#G9281) followed by IRDye 800CW-labeled goat anti-rabbit secondary antibody (LI-COR Cat #926-32211) or IRDye 680-labeled goat anti-mouse secondary antibody (LI-COR Cat #926-68070). Protein bands were visualized using an Odyssey Infrared Imaging System (LI-COR).

To assess virus infectivity, TZM-bl cells (6 × $10^3$ cells/well in 96-well plate seeded one day prior) were challenged with p24-normalized virus via spinoculation (1200 × g, 1 hour, 15°C) in the presence of polybrene (10 μg/ml; Sigma). After spinoculation, the cells were incubated at 37°C and luciferase activity was measured 48 hours after infection using the britelite plus reporter gene assay system (Revvity).

Fluorescent virus particles were analyzed by single virion analysis (23). Virus particles were centrifuged (1200 × g for 30 minutes) onto glass-bottom μ-slides (Ibidi) and then imaged using confocal microscopy. GFP and HALO(JF646) signals in each image were detected using Localize (24) and the percentage of GFP signals that colocalized with JF646 signal was determined using a custom MATLAB (Mathworks) program.

For live-cell imaging experiments, HeLa:POM121-HALO cells were seeded onto glass-bottom μ-slides (3 × $10^4$ cells/well) one day prior to infection. Before infection, the cells were incubated with Complete Media containing 40 nM JF549 dye and incubated for 30 minutes at 37°C. The cells were washed twice with Complete Media to remove excess JF549 dye. Next, the cells were challenged with dual-labeled virus via spinoculation (1200 × g for 1 hour at 15°C) in the presence of polybrene (10 μg/ml) and then incubated at 37°C. At approximately 2 hours post-infection, time-lapse imaging was performed using a Nikon Eclipse Ti-E microscope equipped with a Yokogawa CSU-W1 spinning disk unit and a Plan-Apochromat 100× N.A. 1.49 oil objective. Illumination was provided by 488-nm (GFP), 561-nm (JF549), and 640-nm (JF646) lasers. Confocal z-stacks (3 slices at 0.4 μm step interval centered at the equatorial plane) were acquired every minute for 15 minutes across five fields of view. Images were captured using a 565-nm splitter and two ORCA-fusion BT cameras (Hamamatsu) and analyzed with Nikon Elements or

ImageJ (5). Complete Media containing 2X LEN (or DMSO) was added between the first and second frames, yielding a final LEN concentration of 100 nM. The GFP-CA- labeled particles that remained at the nuclear envelope during the entire 15-minute observation period were analyzed. For display purposes, a pixel-averaging filter was applied to the images, and contrast was adjusted.

### SI Figures

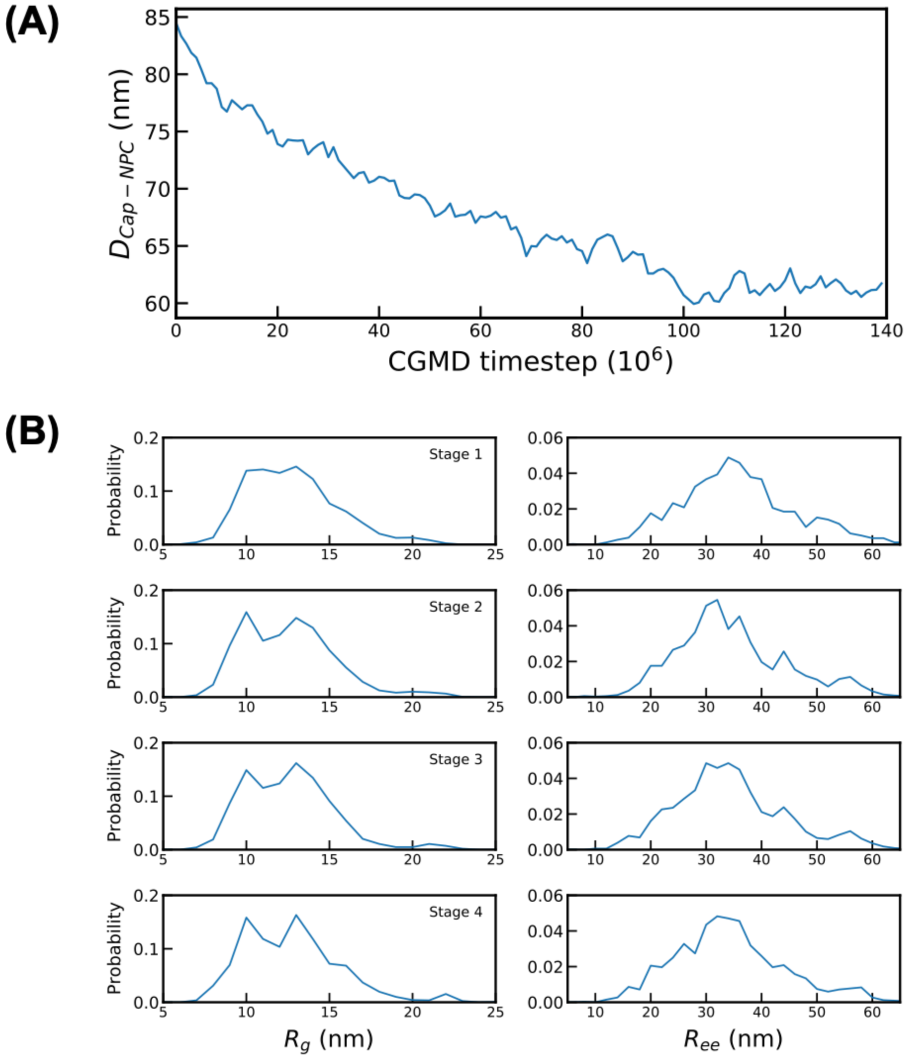


**Figure S1.** Dynamics of capsid translocation (without LEN) to the NPC central channel. (**A**) The time series of the distance ($D_{Cap-NPC}$) between the geometric center of the capsid and equatorial midplane of the NPC inner ring along the channel axis. (**B**) To characterize the conformation of NUP98 chains, we characterize the probability distribution of radius of gyration ($R_g$) and end-to-end distance ($R_{ee}$). Stages 1-4 are defined as equally dividing the simulation time ($140 \times 10^6\, \tau_{CG}$) into 4 equal segments.

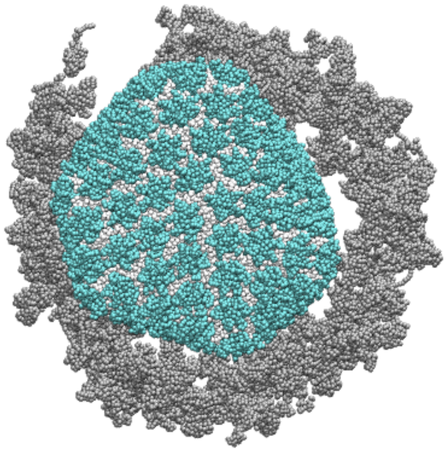

**Figure S2.** Top view of the cross-section of the NPC inner ring when the cone-shaped capsid is docked at the central channel and the NPC is “broken”. The subunits of the inner ring are shown in gray spheres. The NTD domain of both CA hexamer and pentamer are shown in cyan spheres, The CTD domain is shown in white spheres. The rest of the NPC (Y complex, and NUP98 chains), and the nuclear membrane is not shown for clarity.

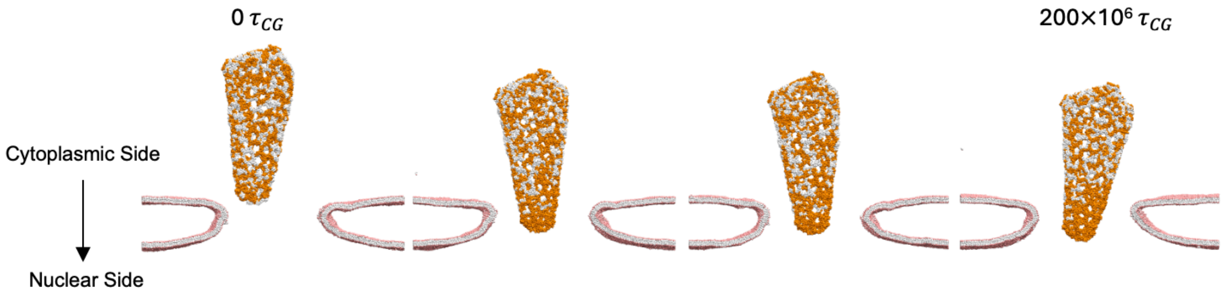


**Figure S3.** Competition between FG-NUP98 and LEN binding during capsid translocation for Replica 2. The capsid is shown in a reduced representation (same as Fig. 2). Only the CTD domain is shown for each CA monomer. The CA monomers to which at least one LEN molecule is bound are shown in white spheres. The CA monomers to which no LEN molecule is bound are shown orange spheres. A cutaway view of the nuclear membrane is also shown to represent the degree of capsid translocation from the cytoplasmic side toward the nuclear side. The NPC, FG-NUP98, and LEN molecules are not shown for clarity.

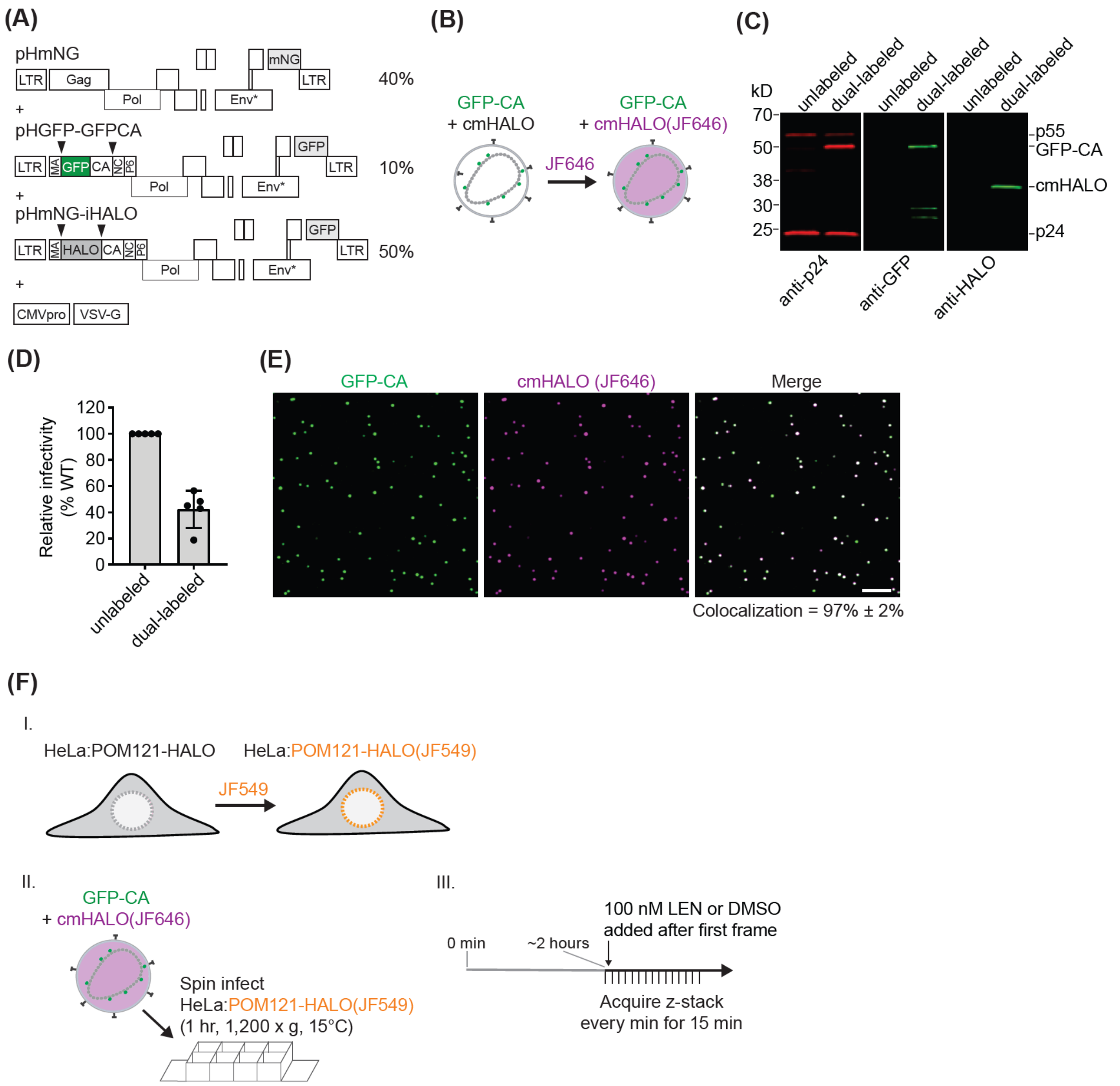


**Figure S4.** Characterization of dual-labeled HIV-1 virions and live-cell imaging assay. **(A)** HIV-1 vector design for virion labeling. HIV-1 plasmids expressing WT Gag (pHmNG) or Gag with internal GFP (pHGFP-GFPCA) or internal HALO (pHmNG-iHALO) were transfected at 40%, 10%, and 50%, respectively, of the total plasmid amount. Black triangles mark protease cleavage sites. Proteolytic cleavage of Gag from pHGFP-GFPCA generates GFP-CA upon virus maturation, while cleavage of Gag from pHmNG-iHALO produces fully processed HALO protein. Asterisk denotes a mutation in the *env* gene introducing a premature stop codon; virions were pseudotyped with VSV-G. The GFP and mNG reporters in the *nef* open reading frame are expressed in virus-producing cells but not incorporated into virions. **(B)** Virions containing cmHALO were labeled with Janelia Farm JF646 dye during transfection. Excess dye was removed during virus concentration. **(C)** Western blot analysis of viral lysates comparing unlabeled and dual-labeled virions. **(D)** Effect of labeling on virus infectivity. TZM-bl cells were infected with p24-normalized amounts of unlabeled or dual-labeled virus. Luciferase activity was measured 48 hours post-infection. **(E)** Representative images of dual-labeled virions. Virions containing GFP-CA and cmHALO(JF646) were centrifuged onto a chambered slide and imaged by confocal microscopy. The percentage of GFP-CA spots that colocalize with cmHALO is shown (AVG ± SD from five images; ~80 virions/image). **(F)** Live-cell imaging of dual-labeled viral cores in HeLa cells stably expressing POM121-HALO. (I) HeLa cells stably expressing POM121-HALO were stained with JF549 dye for 30 minutes, followed by washing to remove excess dye. (II) HeLa:POM121-HALO(JF549) cells were spin-infected with dual-labeled virus. Confocal z-stacks were acquired every minute for 15 minutes beginning ~2 hours post-infection. Complete media containing DMSO or 2X LEN was added between the first and second frames, yielding a final LEN concentration of 100 nM.

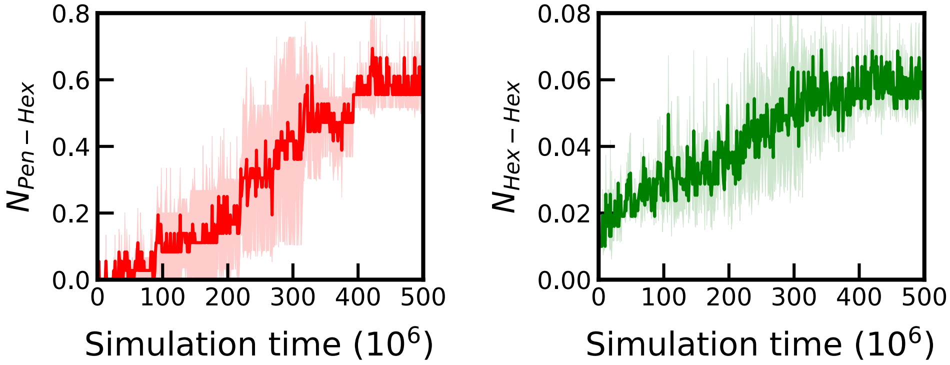


**Figure S5.** Time series of the appearance of defects at the capsid lattice in unbiased CG MD simulations of capsid-LEN complexes in the cytoplasm. The left panel shows the time series of the undercoordinated CA monomers at the hexamer-pentamer interface. The right panel shows the time series of the undercoordinated CA monomers at the hexamer-pentamer interface.

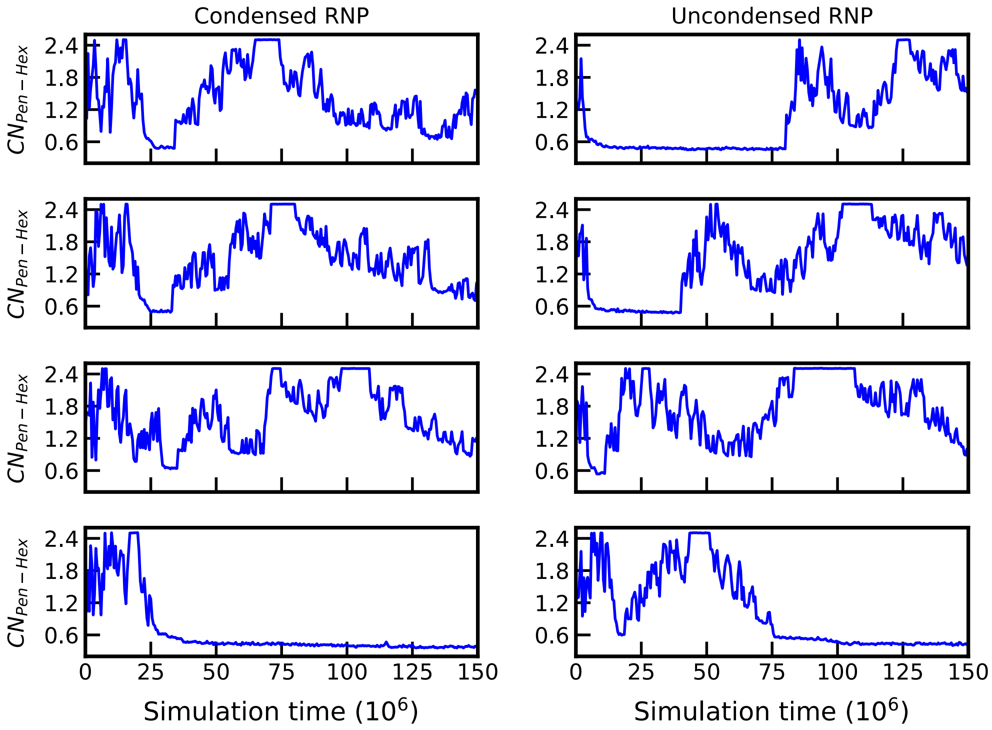


**Figure S6.** Time-series of the collective variable (CV) of WT-MetaD simulations of capsid-LEN complexes in the cytoplasm. 4 replica simulations (from top to bottom) are shown. The CV values fluctuate with time, indicating continuous partial dissociation (which leads to defects) and reformation of contact (healing of defects). In at least one replica (each for both uncondensed RNP and condensed RNP), these defects lead to irreversible rupture of the narrow end within our simulation timescale (CV values less than 0.6).

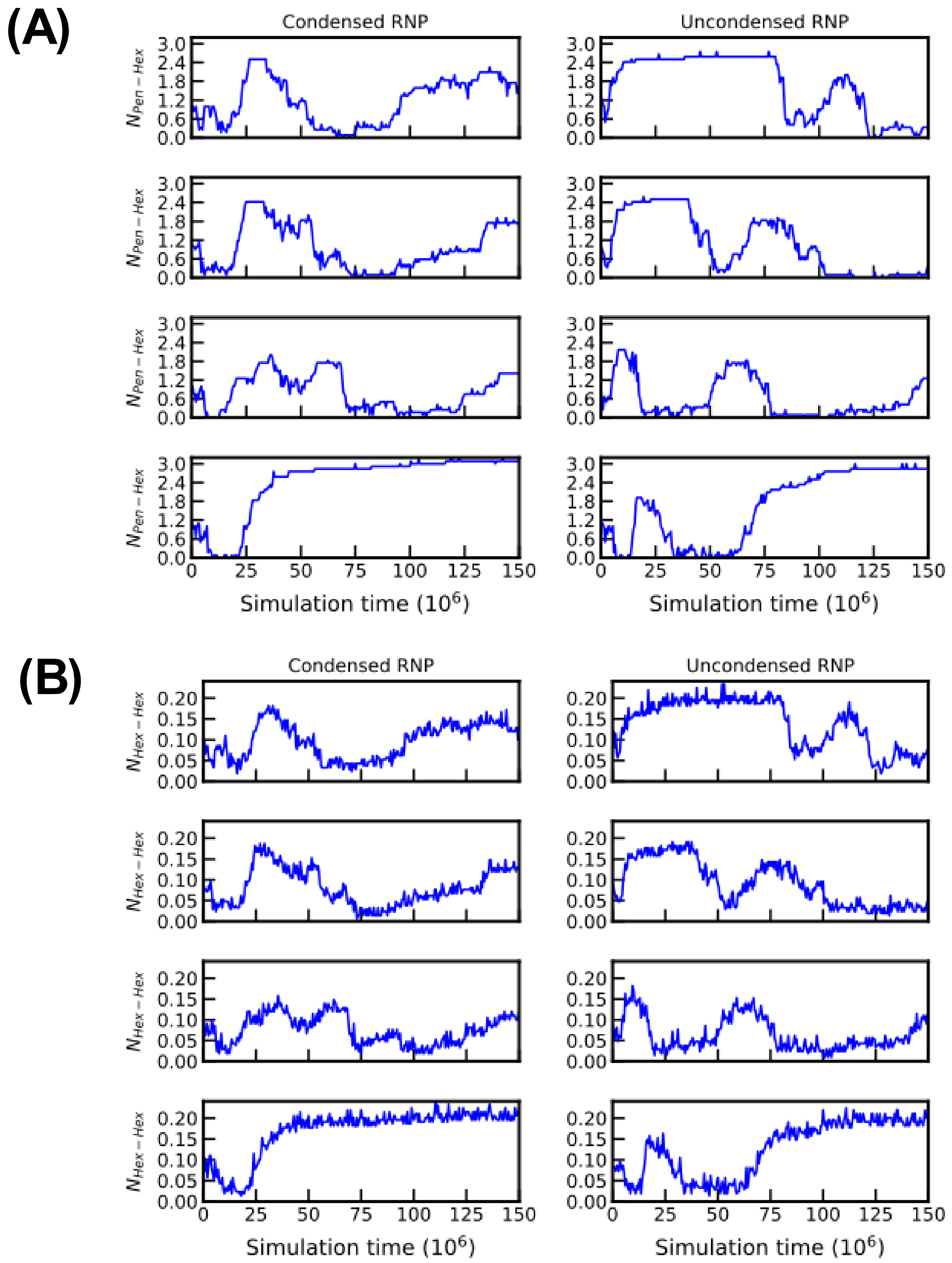


**Figure S7.** Time series of the appearance of defects at the capsid lattice in WT-MetaD simulations of capsid-LEN complexes in the cytoplasm. 4 replica simulations (from top to bottom) are shown. The upper panel (A) shows the time series of the undercoordinated CA monomers at the hexamer-pentamer interface. The lower panel (B) shows the time series of the undercoordinated CA monomers at the hexamer-pentamer interface.

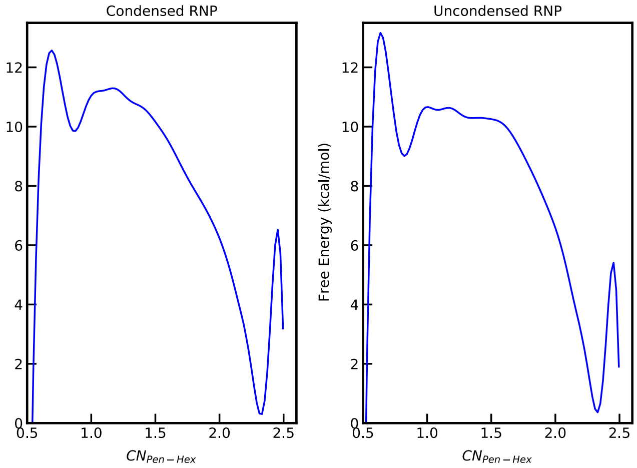


**Figure S8.** Free energy landscape of capsid disassembly of capsid-LEN complexes in the cytoplasm. The units of the free energy are in kcal/mol. The free energy landscape is from cumulative data for all 4 replicas in Fig. S6.

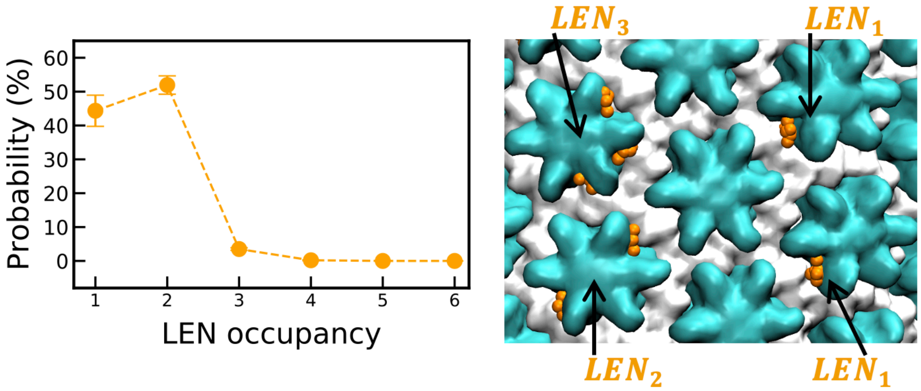


**Figure S9.** Probability distribution of the number of LEN molecules bound to a CA hexamer. The error bars are calculated from 4 replica simulations. The occupancy probability distributions were performed for the final 250 $\times$ $10^6\,\tau_{CG}$. The adjoining snapshot shows 1, 2, and 3 LEN molecules bound to CA hexamer labeled as $LEN_1$, $LEN_2$, and $LEN_3$, respectively.

### SI Movie Legends

**SI Movie 1.** Docking of the cone-shaped capsid (no LEN bound) to the NPC central channel. The NTD domain of CA hexamer and pentamer are shown in cyan and red spheres, respectively. The CTD domain of all CA monomers is shown in white spheres. The NPC is shown in gray, and the nuclear membrane is shown in pink (head group) and white (other groups). The rendered trajectory consists of coordinates saved every 0.5 × $10^6\,\tau_{CG}$ for a cumulative simulation time of 140 × $10^6\,\tau_{CG}$ (Fig. S1). Initially, the NUP98 chains (shown in red) are extended. As the associative interactions between the FG-motifs of NUP98 chains and CA drive the capsid towards the nuclear end from the cytoplasmic end (via an electrostatic "ratchet" mechanism), there is an increasing number of FG-CA interactions that provide the energetic driving force for capsid docking. During the docking trajectory, the NUP98 chains remain in extended conformation which continues to allow extensive FG-CA interactions, thereby providing the energetic driving force for capsid docking to the NPC central channel.

**SI Movie 2 and 3.** Docking of the cone-shaped capsid (LEN-bound) to the NPC central channel (Replica1: SI Movie 2, Replica 2: SI Movie 3). The NTD domain of CA hexamer and pentamer are shown in cyan and red spheres, respectively. The CTD domain of all CA monomers is shown in white spheres. The RNP chains are shown in blue. LEN molecules bound to the capsid are shown in orange. LEN molecules that are not bound to the capsid are not shown for clarity. The NPC (cutaway view) is shown in gray, and the nuclear membrane is shown in pink (head group) and white (other groups). Initially, the CA-CA contacts are dissociated at the hexamer-pentamer interface, first at the narrow end, and then at the wide end. The RNP complex extrudes from these defect sites. As the pentamers are dissociated, the integrity of the capsid is compromised. This leads to the formation of defects at the hexamer-hexamer interface. The movie frames correspond to snapshots every 250000 $\tau_{CG}$.

.

**SI Movie 4 and 5.** Dynamics of LEN-induced rupture of free capsids. The color scheme is the same in other movies. LEN molecules that are not bound to the capsid are not shown for clarity. For ease of viewing, we aligned the capsid coordinates in the movie to the same of the initial frame, which removes the translation degree of freedom. In the SI Movie 4, the RNP is modeled in condensed form. In the SI Movie 5, the RNP is modeled in the uncondensed form.

### SI References:


1. S. Mosalaganti *et al.*, AI-based structure prediction empowers integrative structural analysis of human nuclear pores. *Science* **376**, eabm9506 (2022).
2. Z. Zhang *et al.*, A Systematic Methodology for Defining Coarse-Grained Sites in Large Biomolecules. *Biophysical Journal* **95**, 5073-5083 (2008).
3. M. S. Shell, The relative entropy is fundamental to multiscale and inverse thermodynamic problems. *J. Chem. Phys.* **129**, 144108 (2008).
4. T. C. Beutler, A. E. Mark, R. C. van Schaik, P. R. Gerber, W. F. van Gunsteren, Avoiding singularities and numerical instabilities in free energy calculations based on molecular simulations. *Chemical Physics Letters* **222**, 529-539 (1994).
5. A. Hudait, G. A. Voth, HIV-1 capsid shape, orientation, and entropic elasticity regulate translocation into the nuclear pore complex. *Proceedings of the National Academy of Sciences* **121**, e2313737121 (2024).
6. J. M. A. Grime, J. J. Madsen, Efficient Simulation of Tunable Lipid Assemblies Across Scales and Resolutions. *arXiv e-prints*, arXiv:1910.05362 (2019).
7. S. M. Bester *et al.*, Structural and mechanistic bases for a potent HIV-1 capsid inhibitor. *Science* **370**, 360-364 (2020).
8. C. M. Highland, A. Tan, C. L. Ricaña, J. A. G. Briggs, R. A. Dick, Structural insights into HIV-1 polyanion-dependent capsid lattice formation revealed by single particle cryo-EM. *Proceedings of the National Academy of Sciences* **120**, e2220545120 (2023).
9. S. Plimpton, Fast Parallel Algorithms for Short-Range Molecular Dynamics. *Journal of Computational Physics* **117**, 1-19 (1995).
10. T. Schneider, E. Stoll, Molecular-dynamics study of a three-dimensional one-component model for distortive phase transitions. *Physical Review B* **17**, 1302-1322 (1978).

11. G. J. Martyna, D. J. Tobias, M. L. Klein, Constant pressure molecular dynamics algorithms. *The Journal of Chemical Physics* **101**, 4177-4189 (1994).
12. G. J. Martyna, M. L. Klein, M. Tuckerman, Nosé–Hoover chains: The canonical ensemble via continuous dynamics. *The Journal of Chemical Physics* **97**, 2635-2643 (1992).
13. G. J. Martyna, M. E. Tuckerman, D. J. Tobias, M. L. Klein, Explicit reversible integrators for extended systems dynamics. *Molecular Physics* **87**, 1117-1157 (1996).
14. A. Barducci, G. Bussi, M. Parrinello, Well-Tempered Metadynamics: A Smoothly Converging and Tunable Free-Energy Method. *Physical Review Letters* **100**, 020603 (2008).
15. W. Humphrey, A. Dalke, K. Schulten, VMD: Visual molecular dynamics. *Journal of Molecular Graphics* **14**, 33-38 (1996).
16. W. Lechner, C. Dellago, Accurate determination of crystal structures based on averaged local bond order parameters. *The Journal of Chemical Physics* **129**, 114707 (2008).
17. P. J. Steinhardt, D. R. Nelson, M. Ronchetti, Bond-orientational order in liquids and glasses. *Physical Review B* **28**, 784-805 (1983).
18. E. Lyman, J. Pfaendtner, G. A. Voth, Systematic Multiscale Parameterization of Heterogeneous Elastic Network Models of Proteins. *Biophys. J.* **95**, 4183-4192 (2008).
19. R. C. Burdick *et al.*, Dynamics and regulation of nuclear import and nuclear movements of HIV-1 complexes. *PLOS Pathogens* **13**, e1006570 (2017).
20. J. K. Yee *et al.*, A general method for the generation of high-titer, pantropic retroviral vectors: highly efficient infection of primary hepatocytes. *Proc Natl Acad Sci U S A* **91**, 9564-9568 (1994).
21. R. C. Burdick *et al.*, HIV-1 uncoating requires long double-stranded reverse transcription products. *Science Advances* **10**, eadn7033 (2024).
22. R. C. Burdick *et al.*, HIV-1 uncoats in the nucleus near sites of integration. *Proceedings of the National Academy of Sciences* **117**, 5486-5493 (2020).
23. A. Duchon, R. C. Burdick, V. K. Pathak, W.-S. Hu, "Single-Virion Analysis: A Method to Visualize HIV-1 Particle Content Using Fluorescence Microscopy" in HIV Protocols, V. R. Prasad, G. V. Kalpana, Eds. (Springer US, New York, NY, 2024), 10.1007/978-1-0716-3862-0_6, pp. 77-91.
24. D. Zenklusen, D. R. Larson, R. H. Singer, Single-RNA counting reveals alternative modes of gene expression in yeast. *Nature Structural & Molecular Biology* **15**, 1263-1271 (2008).
